# Dysregulated RNA splicing induces regeneration failure in alcohol-associated liver disease

**DOI:** 10.1101/2024.11.29.626099

**Authors:** Ullas V. Chembazhi, Sushant Bangru, Rajesh Dutta, Diptatanu Das, Brandon Peiffer, Subhashis Natua, Katelyn Toohill, Aurelia Leona, Ishita Purwar, Anuprova Bhowmik, Yogesh Goyal, Zhaoli Sun, Anna Mae Diehl, Auinash Kalsotra

## Abstract

Individuals with progressive liver failure are at a high risk of mortality without liver transplantation. However, our understanding of derailed regenerative responses in failing livers is limited. Here, we performed comprehensive multi-omic profiling of healthy and diseased human livers using bulk and single-nucleus RNA-plus ATAC-seq. We report that hepatic immune milieu alterations in alcohol-associated liver disease (ALD) prevent hepatocytes from transitioning to a proliferative progenitor-like state, trapping them into an unproductive intermediate state. We discovered striking changes in RNA binding protein (RBP) expression, particularly ESRP, PTBP, and SR families, that cause misregulation of developmentally controlled RNA splicing in ALD. Our data pinpoint ESRP2 as a pivotal disease-sensitive RBP and support a causal role of its deficiency in ALD pathogenesis. Notably, splicing defects in ESRP2-targets *Tcf4* and *Slk*, amongst others, directly alter their nuclear localization and activities, disrupting WNT and Hippo signaling pathways, which are critical for normal liver regeneration. We demonstrate that changes in stromal cell populations enrich failing ALD livers with TGF-β, which suppresses ESRP2-driven epithelial splicing program and replaces functional parenchyma with quasi-progenitor-like cells lacking liver-specific functions. This unprecedented account of transcriptional and post-transcriptional dysregulation in ALD suggests that targeting misspliced RNAs could improve recovery and serve as biomarkers for poor ALD outcomes.

## INTRODUCTION

Alcohol-associated liver disease (ALD) stands as the predominant cause of chronic liver disease, with its incidence and prevalence rising in the US and globally^1^. Abstinence remains the sole proven cure for ALD, as both the risk and severity of the disease correlate directly with the dose and duration of alcohol exposure. Similar to other drug-induced liver injuries, liver dysfunction in ALD is most severe shortly after toxin exposure and typically normalizes after the noxious agent is eliminated. However, individuals who develop severe alcohol-associated hepatitis (SAH) or alcohol-associated cirrhosis (AC) are at elevated mortality risk from progressive liver failure despite discontinuing alcohol consumption^2^.

Currently, the only life-saving intervention for ALD patients with acute or chronic liver failure is liver transplantation. Developing alternative treatment options requires understanding the mechanisms that derail normal regenerative responses in livers with SAH or AC. Published work demonstrates a robust correlation between the severity of liver dysfunction and the accumulation of progenitor-like, immature hepatocytes in SAH^3,4^, and senescent hepatocytes in AC^5^ and some SAH patients^6^. There is growing consensus that liver epithelial cells (hepatocytes and cholangiocytes) function as facultative stem/progenitor-like cells to regenerate hepatic parenchyma after adult liver injury^7–9^. Thus, liver failure in ALD likely results from dysregulation of mechanisms that orchestrate state transitions in liver epithelial cells.

Hepatocytes and cholangiocytes in healthy adult livers exhibit high phenotypic plasticity and hence, are able to shed and regain metabolic and proliferative activities^7,9–11^. Although many studies have focused on mechanisms that mediate state transitions by changing gene transcription^12^, post-transcriptional mechanisms that regulate RNA processing, stability, and translation, as well as modulate protein sequence and activity, have proven to be equally critical^13–17^. Post-transcriptional gene regulation is particularly important for implementing rapid responses, during reversible state changes, that liver cells require to survive and reconstitute healthy parenchyma after injury^18–25^. However, the post-transcriptional landscapes of alcohol-associated human liver diseases remain unexplored.

Herein we leveraged next generation deep sequencing of liver RNA and multi-omics platforms (snRNA-seq and snATAC-seq) to identify mechanisms that drive phenotypic differences in liver epithelial cell populations between healthy human livers and explanted human livers with SAH or AC. Putative mediators of the dysregulation were then manipulated in cultured cells and mouse ALD models to determine their pathogenic significance. Our results reveal that RNA processing is globally dysregulated in hepatocytes of livers with decompensated SAH or AC, demonstrate why this occurs, and how it promotes liver failure. These discoveries have important clinical implications as they suggest novel prognostic biomarkers and therapeutic targets for ALD.

## RESULTS

### Altered immune milieu in ALD promotes a quasi-progenitor state of hepatocytes

Recent single-cell resolution studies have significantly advanced our understanding of liver diseases and regenerative processes^11,26–30^. For ALD particularly, these have been hindered by the logistics of tissue preservation and cell isolation for library preparation. To circumvent these challenges, we utilized a 10X multi-omics platform to obtain matched snRNA-seq and snATAC-seq profiles from fresh frozen livers of unaffected individuals, and SAH and AC patients (**Figure 1a**). In parallel, we performed Illumina-based bulk RNA-seq analyses from matched samples to accurately profile lowly expressed genes and post-transcriptional processing. After post-sequencing quality control and analysis (CellRanger ARC, Seurat v4, and Signac, see methods), multi-omic profiling provided 27,000 single nuclei profiles with an average of 1476 genes detected per nucleus. Within this, ∼7000-8000 nuclei were captured from SAH and AC patient livers each, and ∼11,900 nuclei from unaffected samples. All samples were integrated and corrected for batch effects using Harmony. To group and identify clusters, WNN analysis was performed with consideration for gene expression and chromatin accessibility (**Figure 1b**). Cell types were annotated based on differential gene expression and published references for cell-type specific markers^31^. Hepatocytes and non-parenchymal cells (NPCs) clusters were annotated based on expression levels and accessibility of HNF4α and Vimentin (VIM) genes, respectively **(Supplementary Figure 1a**). Within the NPC cluster, we identified most of the major human non-hepatocyte liver cell populations, other than cholangiocytes (**Figure 1b**).

**Figure 1:**
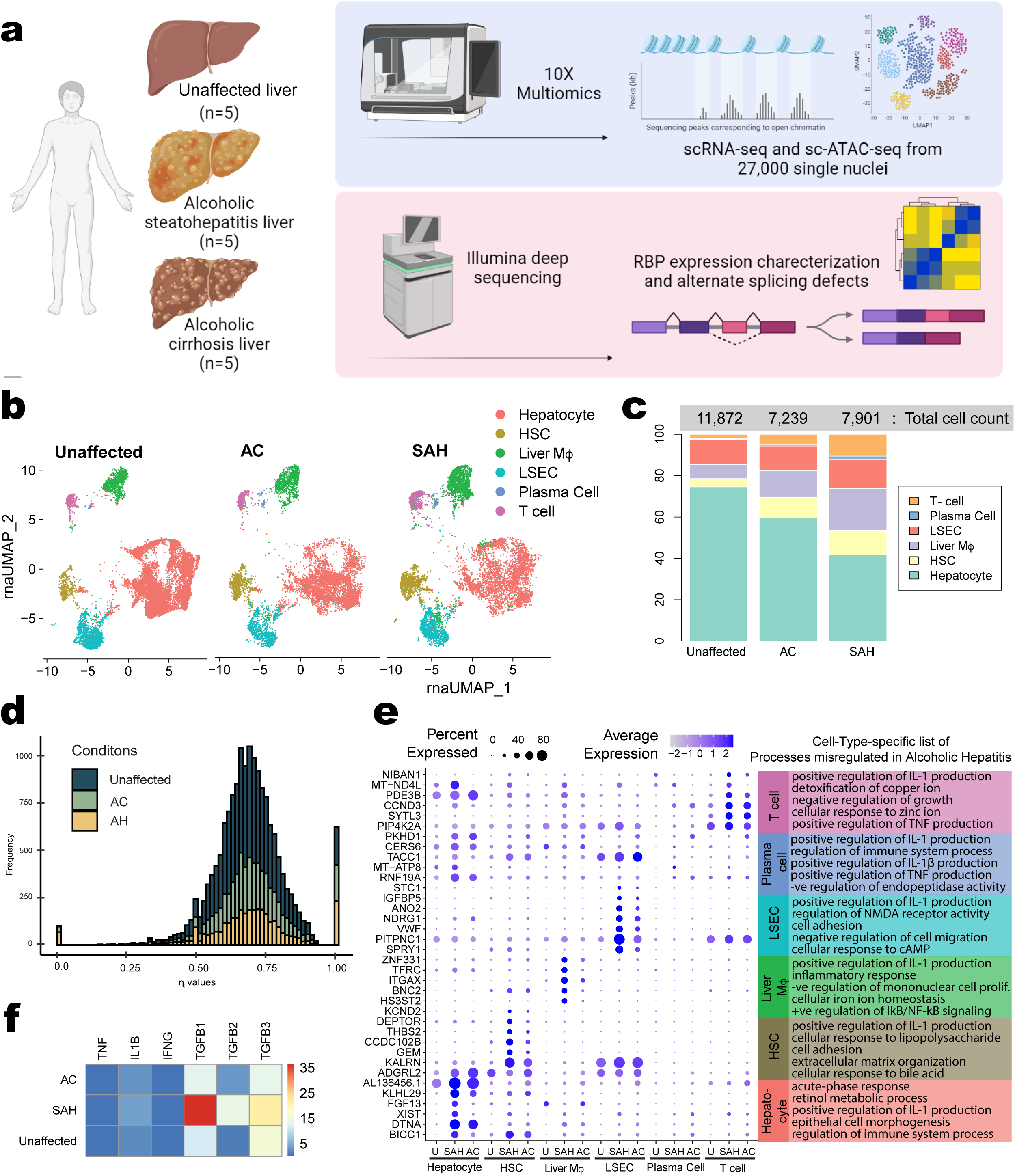
Multi-omic profiling of human alcohol-associated liver diseases. **(a)** Schematic summarizing single nuclear isolation from human liver samples and transcriptomics using multi-omic and deep sequencing approaches **(b)** rnaUMAP showing distinct cell populations identified from multi-omic profiling at single-cell resolution from over 27,000 liver nuclei from 15 individuals (n=5 each) **(c)** Distribution of various liver cells in normal and alcohol-associated liver disease (ALD) patients. AC and SAH showed an increased number of non-hepatocyte (NPC) cells. **(d)** Distribution of porto-central coordinate *η_i_* values in normal liver, AC and SAH conditions. The histogram shows the frequency of cells at different *η_i_* values for each condition. **(e)** Cell type-specific changes in gene expression associated with ALD (examples shown in dot plot, left) and corresponding gene ontology analysis demonstrating affected processes (right). **(f)** Transcript per million (TPM) values of key pro-inflammatory cytokines as quantified from bulk RNA-seq.

Increased inflammation, induced by several underlying factors, is one of the fundamental processes associated with the progression of ALDs. Metabolites, including acetaldehyde, acetate, and reactive oxygen species, produced during the hepatic breakdown of alcohol can induce inflammatory responses. These can be worsened by gut leakiness seen among ALD patients which leads to infiltration of LPS and other microbial products into the liver. Further, ALDs sensitize non-immune cells including hepatocytes to immune signals affecting their normal function. We observed a predominant increase in the fraction of NPCs in diseased samples, led by a 4.5-fold increase in T-cell numbers, indicating widespread inflammation (**Figure 1c**). While liver macrophages showed disease-related increase; we detected a significant shift from the Kupffer cell predominant population to monocyte-derived macrophages in SAH **(Supplementary Figure 1b**). Furthermore, the distribution of hepatocytes along the porto-central axis (see Methods), were affected in both SAH and AC conditions (**Figure 1d, Supplementary Figure 2a, b)**. Specifically, the transcripts expressed in the Portal (zone 1), Midlobular (zone 2), and Central (zone 3) zones exhibited significantly more heterogeneity for SAH and AC than control, as measured by pairwise Euclidian distance in high dimensional PC space **(Supplementary Figure 2c**).

We investigated gene expression programs in SAH and AC patients across various cell populations of the liver (**Figure 1e).** We noticed that differentially expressed genes were enriched in mediators of inflammation-associated signaling pathways such as IL-1, TNF and NFkB. Our analysis indicated that Liver Mβ are the major source of IL1-β, which increased significantly in SAH patients compared to unaffected individuals **(Supplementary Figure 3**). Strikingly, the highest expression levels were observed in Liver Mβ from Cirrhotic livers. In line with previous observations that activated T cells are the major source of immunomodulatory cytokine IFNɣ^32^, we noticed that SAH livers had higher IFNɣ secretion from T cells in parallel to a decrease in lymphocyte-derived TNF-α **(Supplementary Figure 3**). Comparing our findings to bulk RNA-seq data showed that at the transcript level, average levels of TNF-α and IL1-β undergo modest increase, while TGF-β displayed the most striking upregulation in SAH patients on average (**Figure 1f**).

Increased levels of inflammatory cytokines in SAH livers are known to suppress activities of transcription factors that inhibit the proliferation of mature hepatocytes and promote hepatocyte differentiation^33,34^ and this leads to the accumulation of progenitor-like hepatocytes that could be detrimental to SAH patient survival ^3,35^. Hence, we evaluated the activities of transcription factors (TFs) that function to establish hepatocyte identity early in liver development and compared them to changes in TFs that maintain adult hepatocyte identity and function. We first used our matched ATAC-seq dataset from hepatocytes to investigate the activity of ‘adult-specific’ TFs by estimating the enrichment of Tn5 transposase insertions around known motifs corresponding to these TFs (**Figure 2a, 2c, 2d, Supplementary Figure 4a, 4c, 4d)**. This indicated a metagene level decrease in the chromatin accessibility around these adult-specific TF binding sites in AC and SAH patient hepatocytes relative to healthy hepatocytes. In contrast to this, we did not observe a complementary change in the activities of TFs like SOX9 that are active in developing livers, despite an increase in their expression (**Figure 2b, 2c, 2d, Supplementary Figure 4b, 4e, 4f)**. Further, several fetal TFs whose activity is re-established during successful regeneration in mice (e.g., RELA)^11^, did not display a similar activation in ALD hepatocytes (**Figure 2b, Supplementary Figure 4b**). Hence, we propose that the progression of ALDs is associated with a loss of adult-like hepatocyte TF activities, but these livers fail to mount a successful regenerative response due to an incomplete transition to a productive progenitor-like state which normally aids hepatocyte proliferation in response to liver injury^11^.

**Figure 2:**
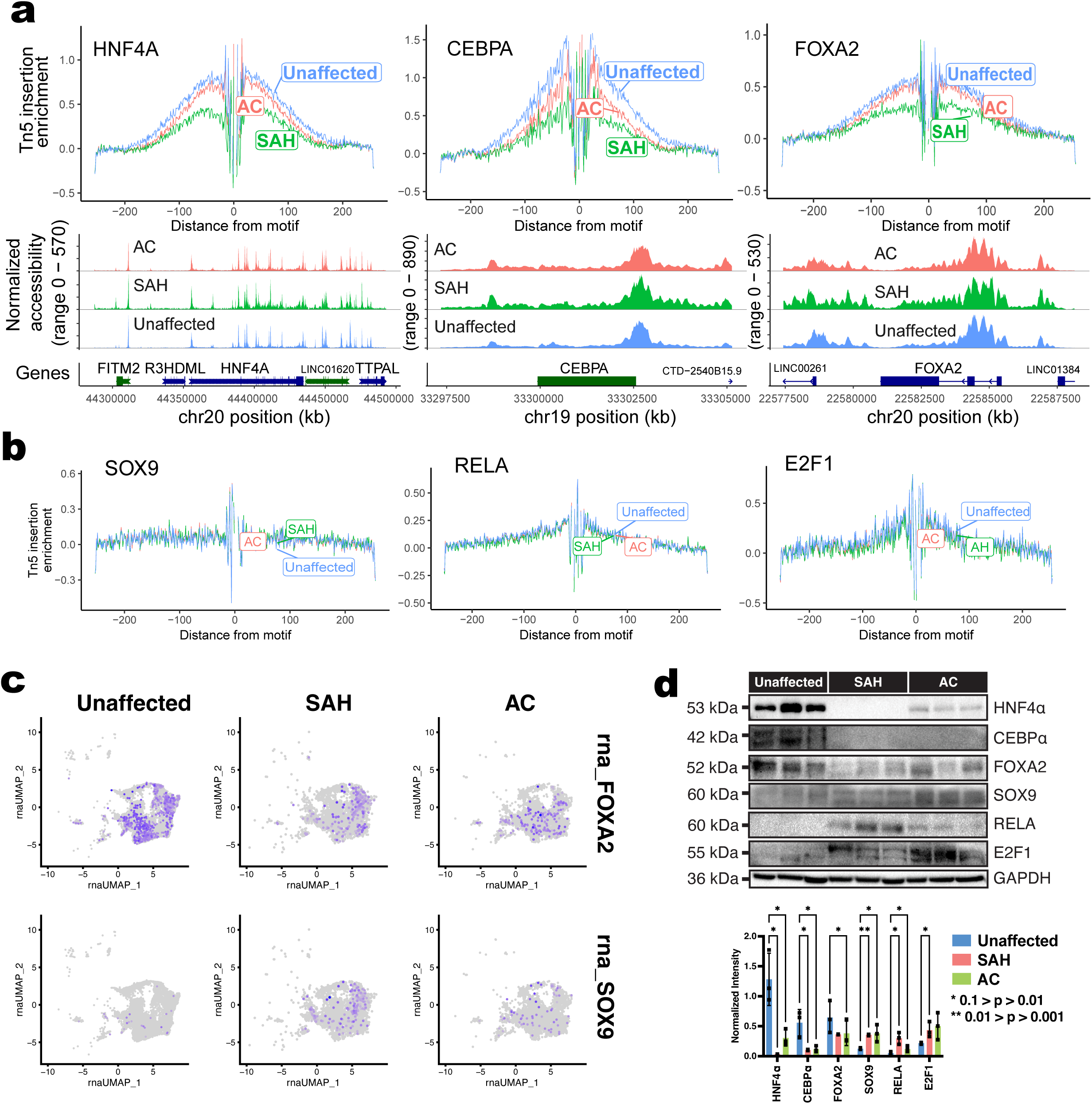
Altered immune milieu in ALD promotes a quasi-progenitor state of hepatocytes. **(a)** (top) Genome-wide Tn5 insertion enrichment plots around motifs for critical transcription factors regulating mature hepatocyte states—HNF4α, CEBPα and FOXA2— showing their decreased activity in SAH and AC patients. (bottom) Genome accessibility around their respective chromatin loci is shown below each plot. **(b)** Genome-wide Tn5 insertion enrichment plots around motifs for critical transcription factors regulating fetal-like hepatocyte states—SOX9, RELA and E2F1—showing their unchanged activity in ALD patients. Chromatin accessibility from snATAC-seq was assessed through Seurat and Signac pipelines (see methods). **(c)** Feature plots showing expression levels of FOXA2 and SOX9 TFs from snRNA-seq. **(d)** Western blot images (above) and corresponding quantitations (below) showing expression levels of TF proteins from human liver lysates (n=3 liver samples per condition).

### Dysregulated RNA binding protein expression and alternative splicing changes in ALDs

We noticed a large fraction of transcripts whose expression levels did not match the chromatin accessibilities at their loci (**Figure 3a, 3b, 3c)**. Such discordance between gene expression and chromatin accessibility is widespread in genomes^36^ and several established examples exist where transcription factors promote gene expression without altering chromatin accessibilities^37–39^. However, alternate mechanisms that operate downstream of transcription could also be driving some of the gene expression changes that occur in diseased liver cells. RNA Binding Proteins (RBPs) orchestrate coordinated changes in groups of RNAs that encode functionally related proteins when cells undergo state transitions^40–42^ and, thus, are attractive candidates as regulators of the dynamic cellular reprogramming that occurs in alcohol-injured livers. Therefore, we used our bulk RNA-seq data to profile human ALD-related differences in RBP expression. We found striking variations in mRNA levels of ∼50 RBPs in diseased versus healthy livers (**Figure 3d, Supplementary Figure 5a**).

**Figure 3:**
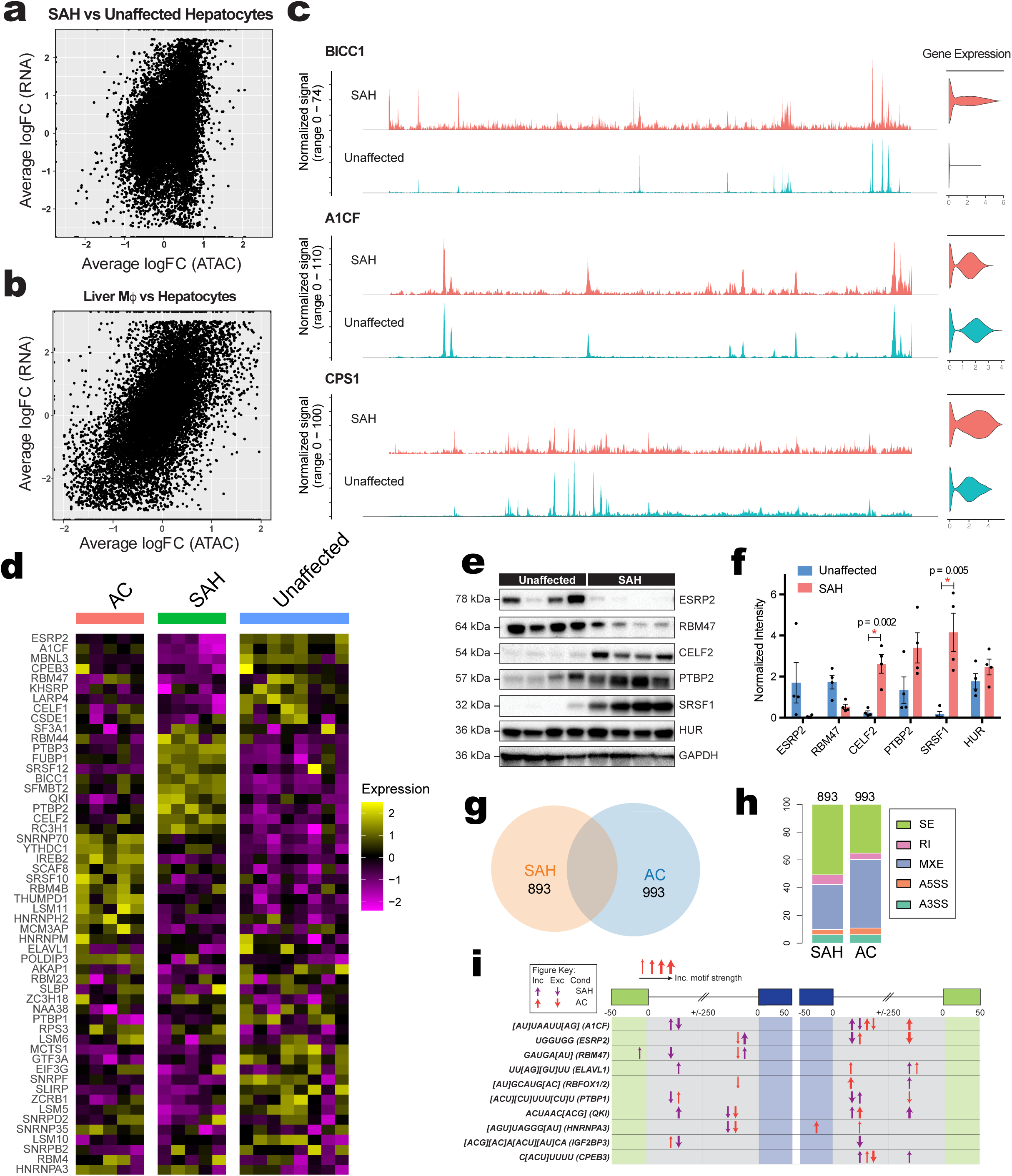
Altered expression levels of RNA binding proteins in SAH and AC induce unique alternative splicing changes. Scatter plot showing a correlation between gene expression and gene accessibility in **(a)** SAH vs unaffected hepatocytes and **(b)** Hepatocytes vs Kupffer cells, indicating that gene accessibility changes strikingly correlate with cell type-specific gene expression changes but not with pathogenic cell-state transitions. **(c)** Examples of concordance and discordance between gene accessibility and expression between SAH and unaffected hepatocytes. **(d)** Heatmap showing expression of top enriched RBPs across unaffected, SAH and AC patients. **(e)** Venn diagram showing the intersection of misregulated alternative splicing events in SAH and AC **(f)** Distribution of alternative splicing events in SAH and AC patients **(g)** Distribution of RNA binding motifs for developmentally regulated alternative splicing factors around AS events. **(h)** Western blot images and **(i)** corresponding quantitation demonstrating protein expression levels of various developmentally regulated RBPs from human livers.

Of note, many of the differentially expressed RBPs are critical regulators of RNA processing and metabolism. Using snRNA-seq-based evaluation of the cell type-specificity of RBPs we found that each cell type had both a unique RBP expression profile and displayed distinct changes in RBP expression patterns in SAH and AC **(Supplementary Figure 5b**). For instance, RBPs such as ESRP2 and A1CF were hepatocyte-specific and downregulated in SAH, whereas CELF2 was primarily expressed in T cells, and its expression was upregulated in SAH **(Supplementary Figure 5b**). Similarly, PTBP2, which serves critical splicing functions during neuronal maturation, was upregulated in liver sinusoidal endothelial cells (LSECs) but downregulated in plasma cells of SAH patients **(Supplementary Figure 5b**). Importantly, the changes in mRNA abundance of RBPs were strongly recapitulated at steady-state protein levels in SAH (**Figure 3e, 3f)**. Particularly, we noticed a significant reduction in the levels of ESRP2 and RBM47, which typically have high expression in adult livers. In contrast, CELF2 and SRSF1— which show decreased expression during liver development—were upregulated in SAH livers compared to unaffected.

The adult RNA splicing factor, ESRP2, was one of the most downregulated RBPs in the SAH livers. ESRP2 is known to regulate alternative splicing (AS) of ∼20% hepatocyte mRNAs and serves important roles in the genesis and maintenance of the adult transcriptome necessary for supporting the mature phenotype of hepatocytes^20,22^. AS adds to the complexity and function of the liver proteome during development, regeneration, and disease. Further, regulated changes in splicing are central to the establishment of identities of several cell types, including hepatocytes^20^. But how it affects the function and progression of ALD has not been systematically investigated. We found that both SAH and AC livers undergo widespread changes in splicing (**Figure 3g);** and we detected 893 and 993 AS events from SAH and AC patient livers, respectively, most of which belonged to the skipped exon (SE) and mutually exclusive exon (MXE) categories (**Figure 3h**). Inspection of RNA sequences surrounding misspliced exons in SAH and AC revealed strong enrichment/de-enrichment of binding motifs for many developmentally regulated RBPs, including ESRP2 (**Figure 3i**). This indicated that altered expression of developmentally regulated RBPs could be driving the splicing changes in SAH by directly binding to the nearby *cis*-regulatory motifs on transcripts.

### Alternative splicing induces widespread functional aberrations in the SAH proteome

We focused on how alternative splicing contributes to disease progression in SAH. We observed that exons misregulated in SAH were distributed across the open-reading frame (ORF), 5’ and 3’ untranslated regions (UTRs), and many overlapped with start and stop codons (**Figure 4a**). A large number of misregulated skipped exons were found in the ORF, implying that exon inclusion/exclusion could generate protein isoforms with altered amino acid sequences and functions (**Figure 4a**). The inclusion/skipping of exons with lengths as multiples of three maintain the reading frame during translation and while most of these events in SAH maintained the ORF frame, there was a significant fraction (∼40%) of exons whose inclusion altered the reading frame (**Figure 4b**). This especially presents a major change in protein isoforms encoded when the misregulated exon is closer to the start codon. We also identified misregulated AS events that led to nonsense-mediated decay, reducing the steady-state transcript levels (**Figure 4b**).

**Figure 4:**
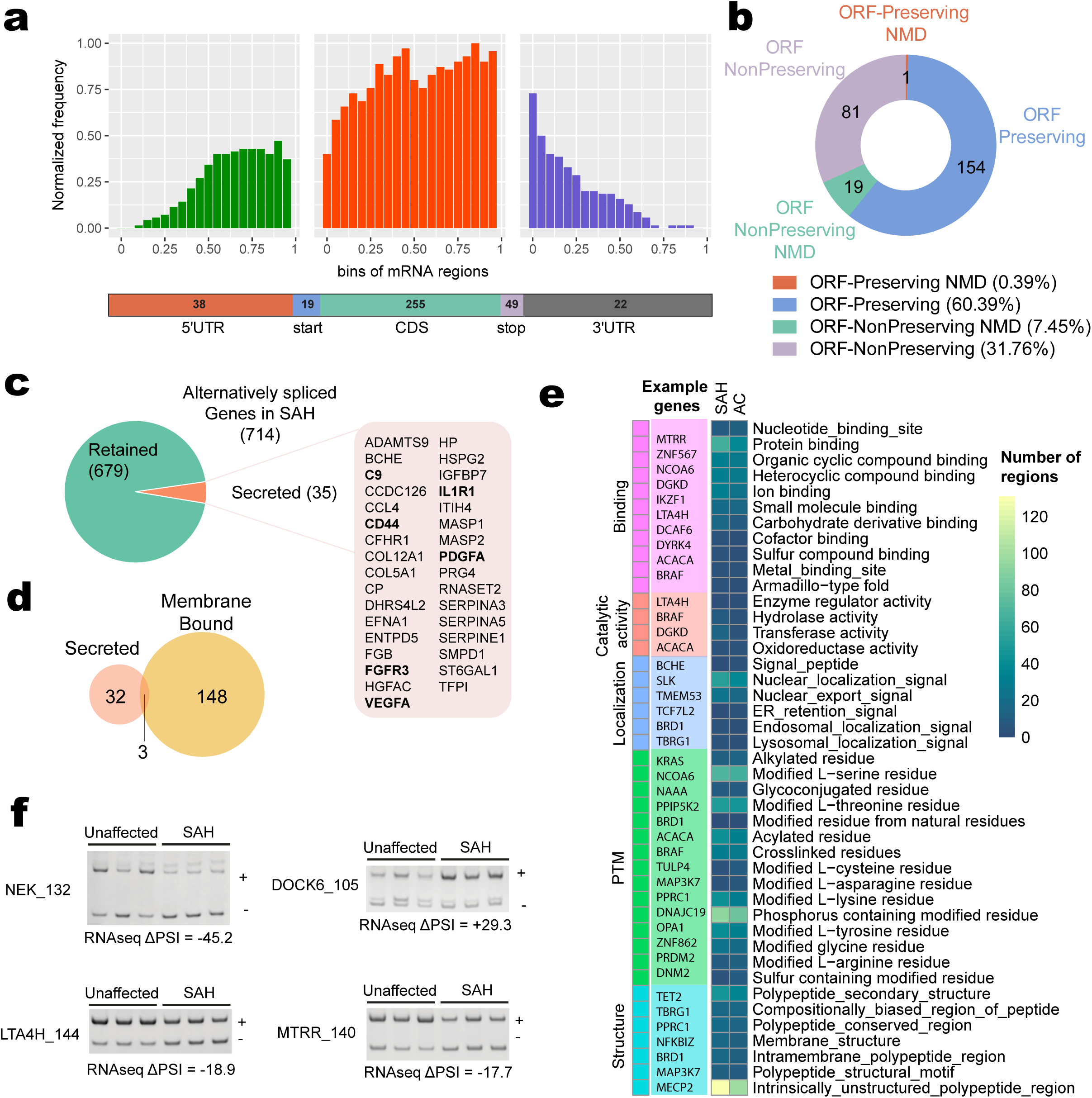
Mis-regulation of alternative splicing causes functional defects in SAH liver proteome. **(a)** Distribution of exons misregulated in SAH patients across gene body and their breakup into transcript regions **(b)** Effect of mis regulated exons on ORF of encoded transcript. **(c)** Pie chart showing the fraction of alternative spliced genes that encode proteins with known secreted isoforms. **(d)** Venn diagram showing the intersection of alternative spliced genes in SAH patients with known secreted and membrane-bound isoforms. **(e)** Heatmap showing numbers of predicted alternative splicing induced functional defects in protein products from SAH and AC patients **(f)** RT-PCR based splice assays validating alternative splicing changes in SAH patients.

Investigating how the inclusion/exclusion of misregulated exons might affect the sequence and function of the encoded proteins, we found that such misspliced transcripts often encoded for secreted proteins. To identify disease-induced alternative protein isoforms among abundant secreted factors, we cross-referenced misspliced genes in SAH to human genes with at least one secreted protein product and found 35 such genes dysregulated in SAH (**Figure 4c**). Out of the 39 misspliced exons identified in these genes, 34 exons had a homology pair in the mouse genome. Notably, we found that the 151 bp exon in FGFR3 and 69 bp exon in PDGFA, which are targets of ESRP2 in mice livers, are misregulated in SAH patient livers. Further, we discovered that of the 714 alternatively spliced genes in SAH, over 20% had membrane-bound protein isoforms, and these included secreted factors FGFR3, CD44, and IL1R1 (**Figure 4d**).

We next explored other functional domains encoded by skipped exons in SAH and AC patients using Exon ontology analysis^43^. This revealed (i) structural features like intrinsically unstructured domains, primary and secondary structure domains, membrane structures, etc. (ii) post-translational modification sites for phosphorylation at serine and threonine residues, alkylation, etc. (iii) intracellular localization signals to nuclei, ER, lysosome, etc. (iv) binding domains for interaction with other proteins, organic cyclic compounds, heterocyclic compounds, etc. and (v) domains critical for the catalytic activity of enzymes such as hydrolases, transferases, etc (**Figure 4e**). We verified many of these individual splicing events using gel-based RT-PCR assays and the results demonstrated excellent correlation with RNA-seq (**Figure 4f**). Despite having fewer misregulated exons than AC, SAH livers showed higher enrichments in several specific categories especially those encoding intrinsically unstructured protein regions, phosphorylation targets, protein binding interactions, and nuclear localization signals.

### Decreased inclusion of NLS containing exons disrupts nuclear localization of SLK and WNT effector TCF4 in SAH livers

We were particularly intrigued by SAH misregulation events that disrupted nuclear localization and nuclear export signals (**Figure 4e**), especially for transcription, and signaling factors, etc. that are critical to hepatocyte functions. We predicted that the exclusion of a nuclear localization domain of a predominantly nuclear protein would lead to its cytoplasmic mislocalization, and negatively influence its function. We evaluated two of such predicted events—93 bp exon in *SLK (SLK_93)* and the 73 bp exon of *TCF4 (TCF4_73)* using RT-PCR-based assays and confirmed that *SLK_93* and *TCF4_73* inclusion decreased by ∼40% and ∼25%, respectively in SAH (**Figure 5a, 5b)**. Immunohistochemistry of SLK and TCF4 proteins revealed that SLK and TCF4 proteins, which are predominantly nuclear localized in unaffected livers, were cytoplasmic localized in SAH patients (**Figure 5c, 5d)**.

**Figure 5:**
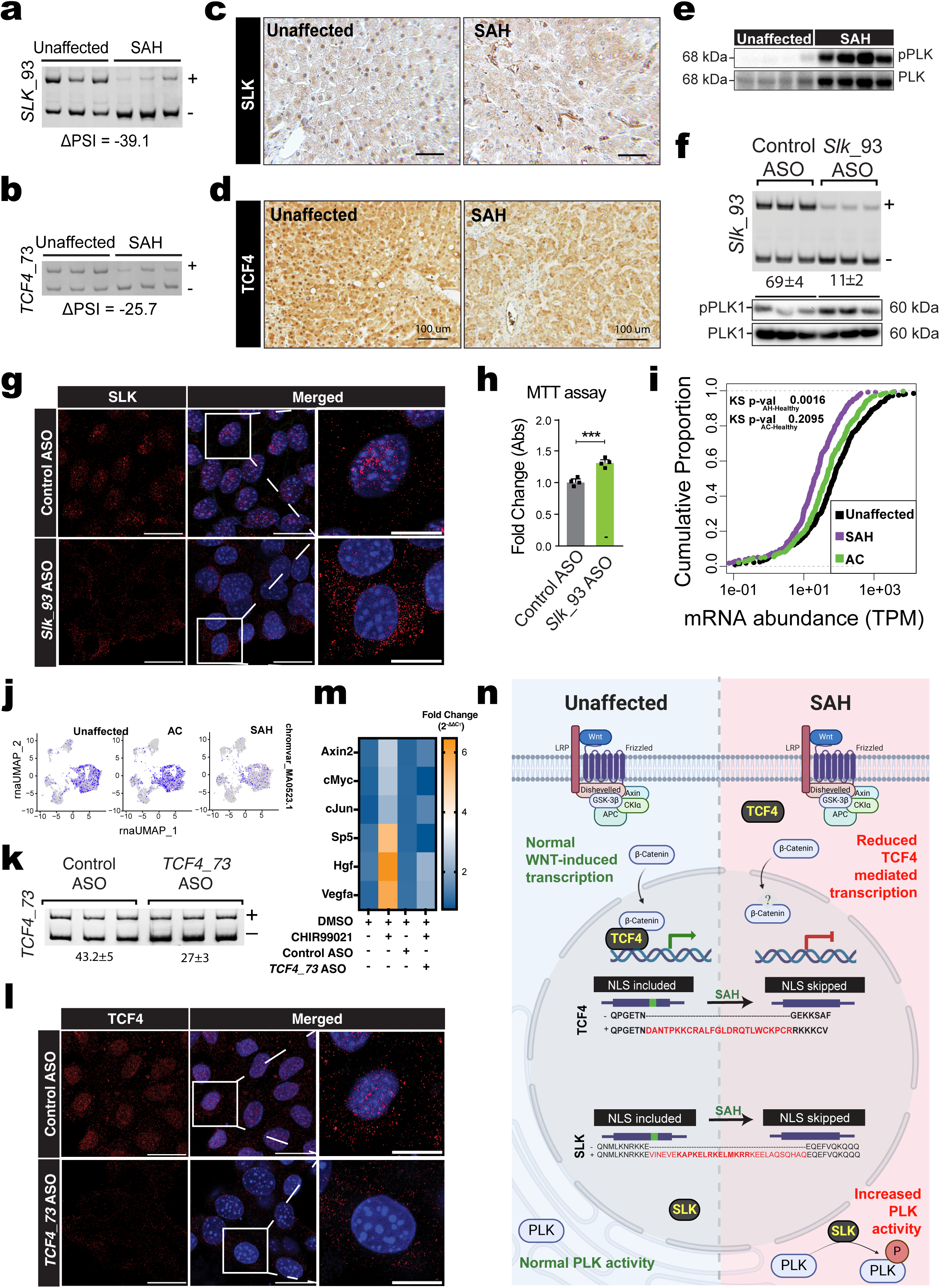
Decreased inclusion of NLS-encoding exon disrupts nuclear localization and function of SLK and TCF4. RT-PCR-based splicing assay demonstrating decreased inclusion of **(a)** 93bp exon of SLK and **(b)** 73 bp exon of TCF4 in SAH patients, both of which encode NLS sequence. IHC images from human patient samples demonstrating that nuclear localization of **(c)** SLK and **(d)** TCF4 is disrupted in SAH patient hepatocytes. **(e)** Western blot analysis from human liver lysates demonstrating increased phosphorylation of Polo-like kinase (PLK) in SAH patients. **(f)** RT-PCR-based splicing assay (top) demonstrating exclusion of the 93 bp exon in SLK transcript upon ASO treatment and western blotting (bottom) showing increased phosphorylation of PLK1 in AML12 cells after ASO treatment. **(g)** Immunofluorescence staining showing decreased nuclear localization of SLK protein in AML12 cells upon ASO-induced exclusion of 93 bp exon in *SLK* transcript. **(h)** quantitation from MTT assay showing increased proliferation in AML12 cells treated with ASO against 93 bp exon of SLK. **(i)** Cumulative plot showing decreased expression of TCF4 target genes in AH patients **(j)** Chromvar plots showing decreased TCF4 activity in SAH hepatocytes in our scATAC-seq dataset**. (k)** RT-PCR-based splicing assay showing increased exclusion of the 73 bp exon in *Tcf4* transcript upon ASO treatment in AML12 cells. **(l)** Immunofluorescence staining showing decreased nuclear localization of SLK protein in AML12 cells upon ASO-induced exclusion of 93 bp exon in SLK transcript. **(m)** Heatmap showing qRT-PCR-based expression levels of WNT downstream target genes in AML12 cells treated with control ASO, CHIR99021 and/or *Tcf4*_73-targeting ASO **(n)** Schematic demonstrating how missplicing in TCF4 and SLK leads to altered WNT signaling and proliferation program in SAH patients.

We next evaluated how these disease-induced AS events lead to functional disruption. Ste20-like kinase (SLK) regulates cytoskeletal modeling and cell migration ^44,45^ and, when in the cytoplasm, it can phosphorylate and activate the multi-stage regulator of mitosis, PLK1 ^46^. We hypothesized that AS-mediated disruption of NLS in SLK protein would lead to its cytoplasmic mislocalization, leading to PLK activation. Western blot (WB) of SLK, PLK, and phosphorylated PLK proteins revealed a relative increase in both the levels of PLK as well as the fraction of phosphorylated PLK in SAH patients, while the SLK levels stayed relatively unchanged (**Figure 5e, Supplementary Figure 6a**).

To test if the AS switch in SLK is sufficient to alter its localization/function, cultured AML12 cells at ∼60-70 % confluency were serum starved and exposed to 1% serum along with *Slk_93*-specific or scrambled control antisense oligo (ASO). The *Slk_93* targeting ASO reduced the inclusion levels to ∼11%, compared to ∼70% inclusion in control ASO treatment (**Figure 5f**). As predicted, immunofluorescence (IF) showed a predominant absence of SLK in the cell nucleus upon ASO treatment (**Figure 5g**), and WB for PLK and phosphorylated PLK revealed significantly increased levels of PLK phosphorylation (**Figure 5f**). Additionally, colorimetric MTT assay revealed a significant increase in absorbance at 550 nm upon treatment with *Slk_93*-targeting ASO, indicating increased proliferation of hepatocytes (**Figure 5h**). These results indicate that expression of cytoplasmic SLK isoform in SAH due to AS promotes PLK phosphorylation and cellular proliferation (**Figure 5n).**

Next, we evaluated TCF4 (also known as TCF7L2) a major co-effector of WNT signaling in the liver. WNT pathway is a critical signaling cascade that is involved in regulating development, homeostasis, regeneration, and disease progression in the liver and controls many aspects of hepatocytes including cell polarity, movement, proliferation, differentiation, zonation, metabolism, and survival^47^. Canonical WNT signaling has β-Catenin as its chief effector, which normally forms a degradation complex with axin, adenomatous polyposis coli (APC), and diversin, facilitating its phosphorylation and degradation. Activation of WNT signaling requires hypo-phosphorylation and release of β-Catenin enabling its nuclear translocation where it partners with T cell factor/lymphoid-enhancing factor (TCF/LEF) family member transcription factor to activate expression of target genes. TCF4 function is key to regulating virtually every aspect of liver function, especially glucose production and other metabolic processes^48,49^. We hypothesized that the cytoplasmic mislocalization caused by decreased inclusion of *TCF4*_73 in SAH patients will disrupt the WNT signaling pathway. Cumulative plot of TCF4 target gene expression revealed a significant decrease in their expression in SAH and AC patients as seen in the leftward shift of these curves compared to the normal individuals (**Figure 5i**). ChromVAR^50^ analysis of snATAC-seq to assess the TCF4 motif activities revealed that unlike normal and AC hepatocytes, SAH hepatocytes had a significant reduction in the chromVAR scores for TCF4 motifs, indicating reduced TCF4 activities (**Figure 5j**). We also found a marked decrease in portal and central genes in SAH compared to the normal livers, suggesting a general loss of zonal identity, and consistent with the role of WNT pathway in driving zonation^47^ **(Supplementary Figure 6b**). As above, using cultured cells we investigated if AS misregulation of *TCF4_73* is sufficient to alter its localization/function. Targeted ASO reduced inclusion levels to 27% (**Figure 5k),** led to a decrease in nuclear-localized TCF4 protein (**Figure 5l**) and abrogated WNT activation by a GSK-3β inhibitor (**Figure 5m**), compared to controls. These data suggest that in SAH, disease-induced alternative splicing causes aberrant localization of TCF4 resulting in a significant decrease in TCF4 activity, especially in zone 3 hepatocytes (**Figure 5n**).

### Loss of ESRP2 exacerbates pathologies associated with extended chronic+binge alcohol diet in mice

To discern RBPs that regulate the splicing of SLK and TCF4 exons, we inspected the genome sequence around *Slk_93* and *Tcf4_73* and identified several occurrences of the 5-mer UUGGG, the consensus *ESRP2* binding motif. Moreover, mice livers with constitutive loss of ESRP2 (ESRP2 KO) showed a significant reduction in the levels of both exons (**Figure 6a, 6b)**. Using *in vivo* eCLIP-seq (enhanced-Crosslinking Immunoprecipitation-seq) of flag-tagged ESRP2, we observed a predominant enrichment of ESRP2 eCLIP Tags (captured binding events) in the upstream intronic region of both the 93 bp *Slk* exon and 73 bp *Tcf4* exon (**Figure 6a, 6b)**, likely indicating direct regulation by ESRP2. Hence, we propose that in SAH patients, a decrease in the levels of ESRP2 induces decreased inclusion of 93 bp exon in *SLK* and 73 bp exon of *TCF4* that encode nuclear localization signals, leading to increased cytoplasmic localization of SLK/TCF4.

**Figure 6:**
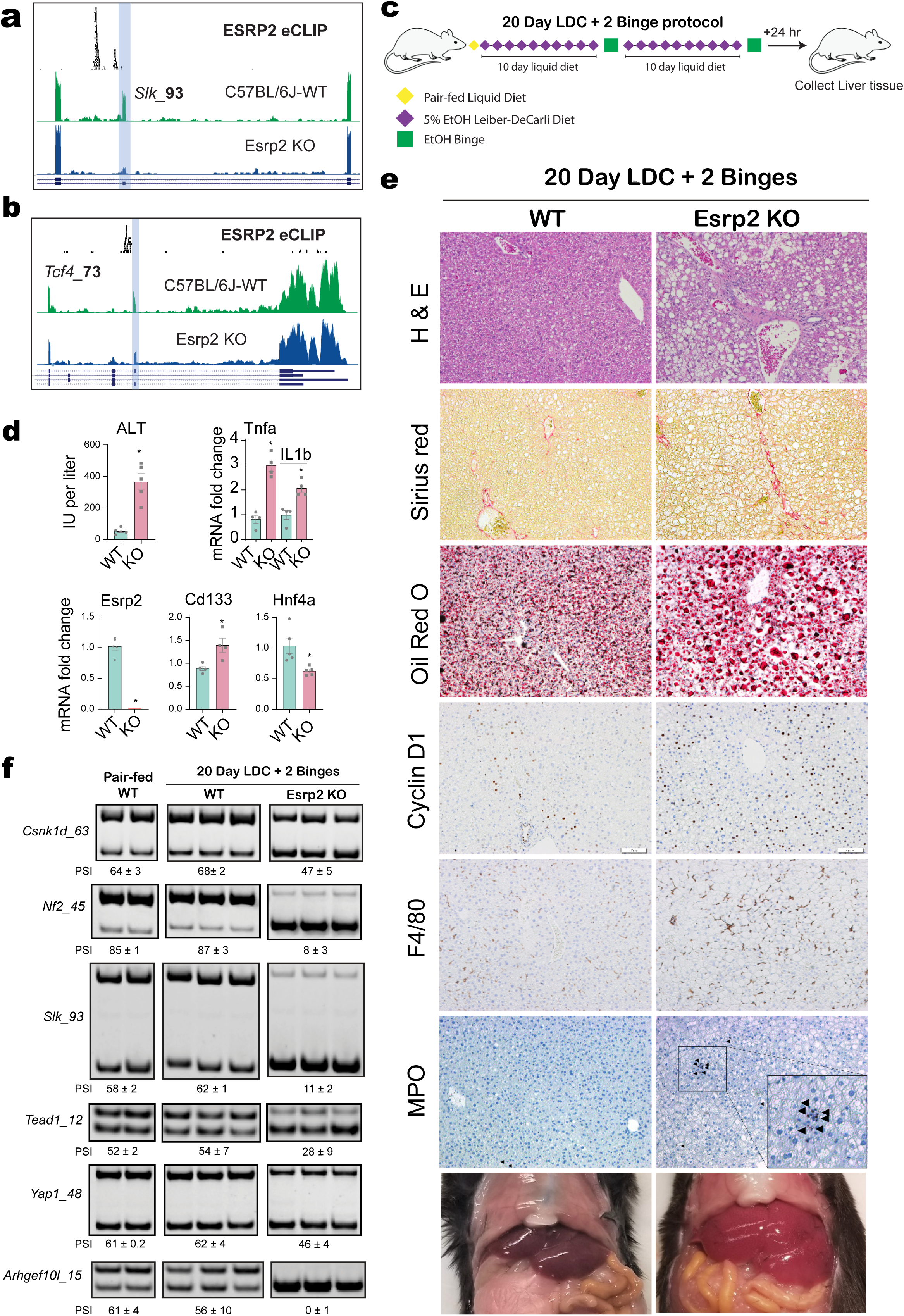
Loss of ESRP2 exacerbates SAH pathology by disrupting developmentally regulated alternative splicing programs. The inclusion of the NLS-containing **(a)** 93 bp *Slk* exon and **(b)** 73 bp *Tcf4* exon decreases significantly in ESRP2 KO mice. ESRP2 binding to the upstream introns of 93 bp *Slk* exon (A) and 73 bp *Tcf4* exon (B) is visualized as eCLIP tags on UCSC genome browser tracks. **(c)** Schematic showing extended NIAAA diet schedule on mice (20D+2B). Mice were fed ad libitum Lieber-DeCarli ethanol liquid diet for 20 days along with two oral gavages of ethanol (5gkg^-1^) – one each at day 10 and day 20. **(d)** ALT levels and qRT-PCR-based quantitation of TNF-α, Esrp2, CD133, and Hnf4a genes. **(e)** Histological characterization of WT and ESRP2 KO mice after 20D+2B diet. **(f)** Gel-based RT-PCR assay showing ESRP2 depletion leads to SAH-like transitions in alternative splicing profiles in 20D+2B mice liver.

As shown above, the consensus motif for ESRP2 was one of the most enriched/de-enriched around the exons that are alternatively spliced in SAH/AC patients, primarily in the upstream/downstream intronic regions **(Supplementary Figure 7, Figure 3c**). We next characterized transcriptome-wide changes associated with alcohol feeding in mice and how ESRP2 mediates the maintenance of adult-like characteristics of hepatocytes in this context. To recapitulate the early stages of alcohol-associated hepatitis, we devised an extended version of the NIAAA model for our studies (20D+2B) where mice were fed *ad libitum* Lieber-DeCarli ethanol liquid diet for 20 days and administered two oral gavages of ethanol (5g/kg^-1^) – one each at day 10 and day 20 (**Figure 6c**). Livers were collected 24 hours after the second binge to profile transcriptomic changes. We found that 20D+2B fed mice recapitulated over 20% of the gene expression defects seen in human SAH patients. Importantly, mice lacking ESRP2 exhibited exacerbated alcohol-diet-associated phenotypes on the 20D+2B diet, in comparison to treated WT mice **(Supplementary Figure 8a, 8b)**. Specifically, ESRP2 KO mice exhibited severe hepatocyte injury, increased levels of inflammatory cytokines, and decreased expression of adult hepatocyte transcription factors (e.g. HNF4α) (**Figure 6d**). Significant increase in steatosis (as seen from H&E and Oil red O staining), inflammation (as seen from MPO and F480 staining), cellular proliferation (as seen from Cyclin D1 staining) were noted, and portal fibrosis was enhanced in the ESRP2 KO mice (**Figure 6e**).

While both WT and KO showed splicing changes in several genes upon 20D+2B alcohol feeding, ESRP2 KO mice recapitulated splicing changes in a larger proportion of genes misregulated in human SAH patients **(Supplementary Figure 8c**). Several of the splicing defects in genes including *Slk, Arhgef10L* and key regulators of the Hippo pathway such as *Nf2*, *Tead1*, *Yap,* etc. were validated by RT-PCR (**Figure 6f**). While alcohol feeding in WT mice only produced modest changes, alcohol-fed ESRP2 KO mice showed a strong misregulation in alternative splicing. This was particularly the case for *SLK_93* in ESRP2 KO mice, which closely resembled levels observed in human SAH livers. Further, in line with our predictions about adult-to-fetal reprogramming responses, ESRP2 KO mice demonstrated increased proliferation on the alcohol diet, which could partially be explained by the expression of the cytoplasmic isoform of *SLK*. Similarly, the cumulative plot of TCF4 target gene expression revealed a significant decrease in their expression in ESRP2 KO mice compared to other control groups, indicating reduced activity of TCF4 in ESRP2 KO mice relative to WT mice on alcohol diet **(Supplementary Figure 8d**).

### Increased TGF-β signaling drives ESRP2 suppression in SAH

To pinpoint upstream signaling events most critical to suppressing the adult identity of hepatocytes in SAH, we implemented the Niche-net pipeline to identify ligands that influence expression in target cells and predict intermediate signaling factors. We started with a gene list of interest, previously benchmarked for predicting the successful differentiation of hepatocytes from iPSCs, that models genes that potentially predict hepatocyte maturity. We performed a data-driven prioritization of ligands expressed in non-parenchymal cell populations based on how well they predict the expression of this gene list of interest (**Figure 7a**). Among the top identified ligands, TGF-β showed increased expression in most non-parenchymal cell populations in SAH livers, with highest expression levels in T lymphocytes (**Figure 7a, 7b)**. Furthermore. TGF-β displayed strong regulatory potential on several target genes, including PPARg, SOX9, CTNNB1, and HIF1A (**Figure 7a**). We also identified HGF as another ligand that could regulate hepatocyte identity by potentially regulating the expression of MET.

**Figure 7:**
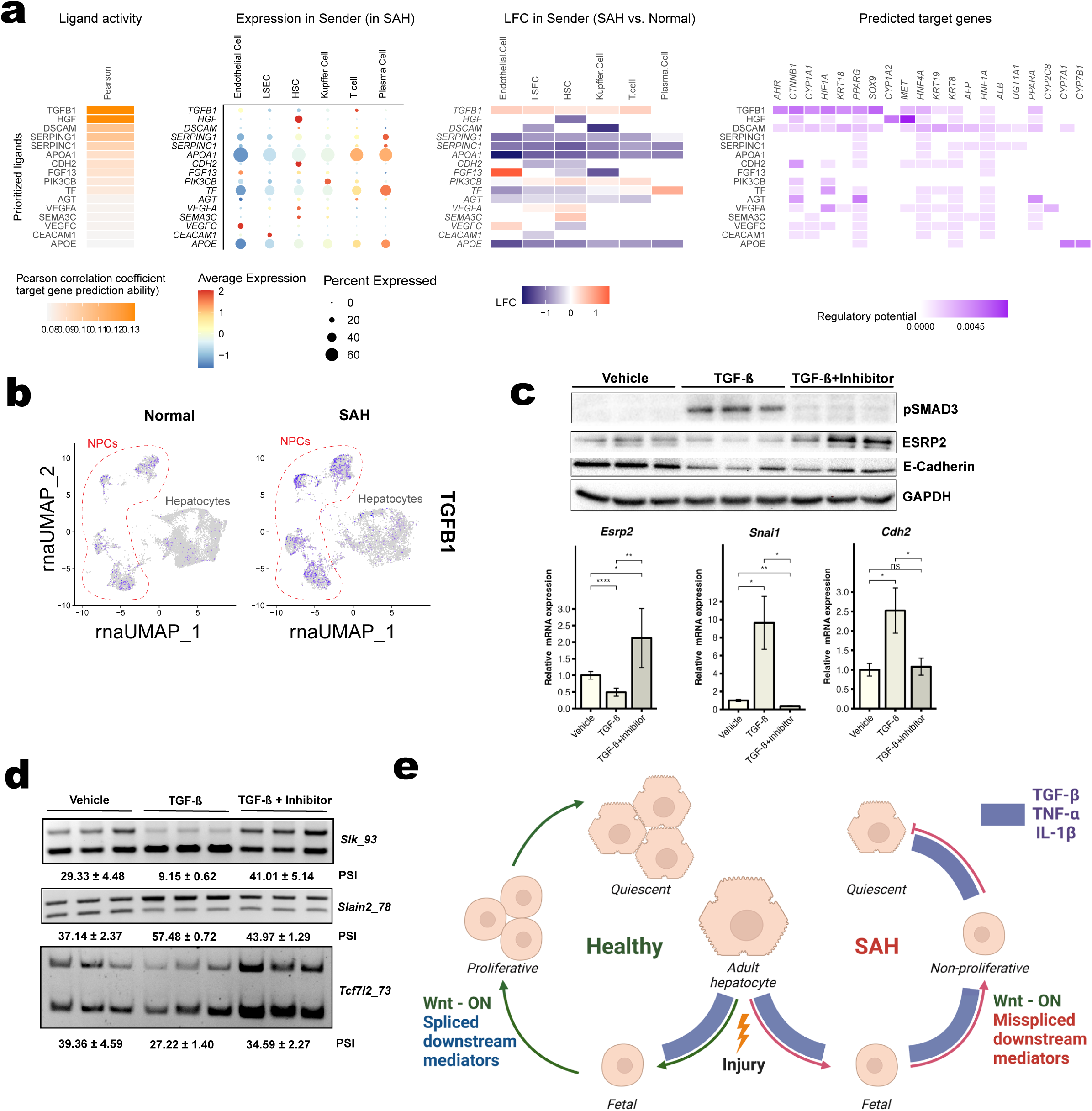
TGF-β signaling in SAH livers is a key determinant of hepatocyte identities. **(a)** Cell interaction analysis summarizing key ligands that regulate hepatocyte identities in SAH livers, log fold changes (LFC) in their expression levels in SAH livers compared to unaffected livers, and their potential to regulate key hepatocyte target genes. TGF-β1 and HGF ligands showed the highest correlation to the gene set defining hepatocyte maturity. **(b)** Feature plot illustrating increased expression levels of TGF-β1 specifically in the NPC populations from SAH patients. **(c)** Western blot and RT-qPCR analyses showing ESRP2 reduction upon TGF-β1 treatment (5 ng/mL in DMEM/F-12 complete media for 36 hours) and restoration upon addition of TGF-βR I/II inhibitor LY2109761 (50 uM for 48 hours). TGF-β1 treatment increases steady-state mRNA levels of EMT-transcription factors *Snai1* and *Cdh2*, which return to baseline after inhibitor addition. Phospho-Smad3 expression confirms TGF-β pathway activation and its subsequent reduction upon inhibitor treatment confirms inhibitor efficacy. **(d)** RT-PCR analysis of ESRP2 splicing targets (*Slk*, *Slain2*, and *Tcf4*) demonstrates significant exon skipping (93 bp, 78 bp, and 73 bp, respectively) following TGF-β treatment. These levels return to baseline upon the addition of the TGF-β inhibitor. Exon lengths are shown beside the gene name, and the differences in PSI values are shown as mean ± SDM below each image. **(e)** Schematic showing a proposed model of the combinatory roles of cytokines and Wnt signaling in regulating cellular transitions in hepatocytes after injury, which is severely misregulated in SAH due to misspliced downstream Wnt mediators.

TGF-β signaling is critical for the initiation and maintenance of epithelial-to-mesenchymal transition (EMT) by upregulating key EMT regulators, including Snail, δEF1/SIP1, and Id proteins. Importantly, δEF1/SIP1, Snail, and Twist bind to the promoter regions of ESRPs to suppress their expression and downstream splicing activities^51,52^. To test the direct contribution of TGF-β in SAH-related downregulation of ESRP2, we treated hepatocyte cultures with TGF-β for 36 hours. Hepatocytes exposed to TGF-β showed significant upregulation of SNAI1 mRNA and increased phosphorylation of SMAD3 protein (**Figure 7c**). Additionally, TGF-β treatment resulted in increased N-cadherin (*Cdh2*) mRNA abundance and a pronounced reduction in ESRP2 and E-Cadherin protein levels (**Figure 7c**). Further, we noted similar changes in the splicing patterns of *Slk* and *Tcf4* mRNAs upon TGF-β treatment (**Figure 7d**) that were initially detected in the SAH livers. To explore the potential recovery of ESRP2 and its splicing targets in ALD, we introduced a TGF-β receptor type I/II inhibitor (LY2109761), which suppresses both basal and TGF-β-stimulated cell migration. Treatment of TGF-β-exposed cells with the inhibitor reduced pSmad3 levels, indicating effective blockade of the TGF-β signaling pathway (**Figure 7c**). Importantly, treating hepatocyte cultures with the inhibitor reinstated ESRP2 expression and restored the splicing of its targets transcripts (*Slk*, *Tcf4*, and *Slain2*) (**Figure 7c, d)**. Together, these findings highlight the therapeutic potential of TGF-β inhibitors in counteracting ALD-associated spliceopathy, hepatocyte dedifferentiation and regeneration failure.

## DISCUSSION

SAH and AC are progressive stages of ALD. SAH is characterized by hepatocyte death and presents with jaundice, hepatic encephalopathy, and portal hypertension. As the disease progresses, chronic inflammation from alcohol-associated hepatitis induces extensive scarring and irreversible liver damage, transforming liver from tender and enlarged in SAH to small and firm in AC. Surviving liver epithelial cells in patients with SAH and decompensated AC are dysfunctional and unable to regenerate sufficient healthy parenchyma to avert liver failure^3,4^. Previous studies have shown that transcriptomes of ALD patient livers undergo significant reprogramming and expression of transcription factors controlling hepatocyte maturation is de-regulated in ALD^53^. In line with these findings, we found that the abundance, nuclear localization, and activities of transcription factors that maintain the mature hepatocyte phenotype (e.g., HNF4α, C/EBPα, FOXA2) are reduced in SAH and AC although transcription of the transcription factors themselves is similarly active in diseased and healthy livers. Further, the activities of transcription factors that enable hepatocytes to transition to a less mature, more proliferative phenotype (e.g., SOX9, RELA, E2F1) are not increased despite disease-related induction of these transcription factors. These findings indicate that mechanisms operating downstream of transcription critically regulate the phenotypic transitions that are required to recover functional hepatic parenchyma in livers that were injured by alcohol. Dysregulation of these processes traps hepatocytes in an intermediate state where they are neither fully mature nor functional progenitors (**Figure 7e**).

We analyzed explanted livers that are enriched with these dysfunctional cells to identify mechanisms that maintain their persistence long after the removal of the toxin that induced the liver damage. Our data show that global changes in the hepatic expression of RBPs significantly contribute to this pathology. RBPs interact with microRNAs and other noncoding RNAs to control the stability, splicing, and translation of suites of mRNAs that must collaborate to orchestrate complex phenotypic switches^54^. Previous studies have demonstrated the importance of the microRNA network in modulating susceptibility to alcohol-related hepatotoxicity^55,56^. Our work presents new evidence about various post-transcriptional mechanisms that dictate ALD outcomes in several ways. The results: *i)* demonstrate disease-related differences in the mRNA expression of a large number of RBPs; *ii)* prove that the differences in transcript abundance are paralleled by differences in protein expression of these RBPs, as well as *iii)* changes in their mRNA targets that *iv)* result in functional changes in the proteins encoded by those mRNAs, and lead to *v)* changes in signaling that broadly control reparative and regenerative responses in injured livers. Together, these foundational findings provide a platform for studies to identify specific RBPs, which function as the hubs that tether this interactome as these RBPs (or some of their targets) could be manipulated to improve recovery from ALD.

We focused on ESRP2 as a pivotal disease-sensitive RBP because it is silenced in the early stages of liver development but *i)* up-regulated dramatically in the final stages of hepatocyte differentiation, *ii)* expressed specifically in hepatocytes of healthy adult livers, and *iii)* known to regulate the adult splicing program of ∼20% of the mature hepatocyte genome^20^. Further, the current analysis confirmed that hepatocyte expression of ESRP2 is virtually undetectable in livers with clinically decompensated ALD^33^. Thus, ESRP2 is one of the most severely downregulated RBPs in the transcriptome and proteome of hepatocytes in livers with SAH, and its expression is also significantly suppressed in AC. ESRP2 prevents hepatocytes from undergoing EMT and thereby maintains hepatic epithelial integrity. Epithelial integrity is essential to execute vital liver-specific functions. After liver injury, cell-to-cell connections in surviving hepatocytes must be relaxed, but then re-established efficiently, for liver regeneration to reconstitute functional hepatic parenchyma. Consistent with this notion, our earlier work showed that injury-related cytokines that initiate liver regeneration (e.g., TNF-α, IL1-β) suppress ESRP2^33^. Downregulation of ESRP2 inhibits Hippo kinase signaling to enable activation of YAP, a factor that promotes hepatocyte EMT^20^. In this study, we showed that decreasing ESRP2 activates Polo-like kinase 1 (PLK-1), which is known to expedite the transition through later stages of the cell cycle and facilitate hepatocyte proliferation^57^. After partial hepatectomy, TNF-α and IL1-β are transiently induced, ESRP2 is transiently repressed, and effective liver regeneration ensues.

When ESRP2 suppression is sustained in hepatocytes of injured livers, however, regeneration is defective and maladaptive inflammatory and fibrogenic repair responses predominate. Our mouse model of ALD supports a causal role for ESRP2 depletion in this liver growth dysregulation by showing that Esrp2 KO mice develop significantly worse liver inflammation, fibrosis, and spliceopathy than alcohol-fed controls. Our single nuclei analysis of human SAH and AC livers (which ‘naturally’ accumulate ESRP2-deficient hepatocytes) demonstrates significant qualitative and quantitative changes in immune cell- and hepatic stellate cell-populations. These changes in stromal cell populations enrich the liver microenvironment with factors that promote inflammation (e.g., TNF-α, IL1-β) and fibrosis (e.g., TGF-β); all of which also suppress ESRP2. Hence, the hepatic microenvironment in ALD favors sustained downregulation of ESRP2 and the resultant persistence of an immature state of hepatocytes.

Recent studies have indicated that adult hepatocytes require three processes to replicate - de-differentiative signaling initiated by pro-inflammatory cytokines, proliferative signaling driven by growth factors/Wnts, and inhibition of TGF-β signaling^58^. Although hepatocytes of alcohol-fed ESRP2 KO mice and humans with SAH or AC switch to a less mature phenotype, they may be growth-arrested given that net hepatic expression of Wnt target genes is suppressed and the hepatocyte populations are relatively depleted. This evidence of reduced regenerative activity is paradoxical because liver size is not reduced in ESRP2 KO mice^20^ and herein we show that depleting ESRP2 in healthy hepatocytes activates PLK1, a response that is predicted to expedite transition through late (i.e., G2-M) phases of the cell cycle. On the other hand, aberrant activation of PLK1 in injured tissues has been reported to subvert responses that normally repair damaged DNA, leading to mitotic catastrophe and cell senescence^57^. Indeed, we observed that alcohol-fed ESRP2 KO mice accumulate cyclin D1-positive hepatocytes in hepatic acinar zone 2, where hepatocytes in pre-replicative phases of the cell cycle accumulate in healthy liver^8,11,59^. Together, these observations suggest that many surviving ESRP2-deficient hepatocytes in livers with SAH or AC may be arrested in pre-replicative stages of the cell cycle and thus, non-regenerative despite exhibiting a progenitor-like phenotype (**Figure 7e**). This possibility merits further study in light of a recent report that numbers of biliary-derived bi-potent transitional liver progenitor cells (TLPCs) increase in parallel with accumulation of senescent hepatocytes in many liver diseases, including ALD. TLPCs must differentiate into hepatocytes before replicating^9^. ESRP2 expression is hepatocyte-restricted in adult liver^22^, raising the possibility that proper regulation of ESRP2 targets is required to regenerate healthy liver tissue after injury. Some of these ESRP2-regulated factors may be novel therapeutic targets; others (e.g., aberrantly spliced secreted proteins) may simply telegraph dysfunctional repair responses and serve as novel biomarkers for poor ALD outcomes.

## METHODS

### Sample Collection

Human liver samples were acquired from patient explants (right lobe) and donor resections (caudate lobe). Collected samples were either snap-frozen and stored at −80°C or formalin-fixed and embedded in paraffin until use. The diagnosis of SAH and AC is based on previously published guidelines^60,61^ and all patients with SAH had some degree of cirrhosis or advanced fibrosis. Human samples were received from the Department of Surgery at Johns Hopkins Hospital (supported by the NIAAA, R24AA025017, Clinical resources for AH investigators). The use of liver samples from patients with SAH, AC, and donor controls was approved by the Institutional Review Board (IRB00107893 and IRB00154881) at Johns Hopkins University.

### Experimental Animals

National Institutes of Health (NIH) and institutional guidelines were followed in the use and care of laboratory animals and experimental protocols were performed as approved by the Institutional Animal Care and Use Committee (IACUC) at University of Illinois, Urbana-Chamapign and Duke University. Mice were housed on a standard 12-hour-light/dark cycle (18-23 °C ambient temperature; 40-60% humidity) and were allowed *ad libitum* access to water and a normal chow diet (2918 Envigo Teklad). Genomic DNA was derived from tail biopsies and genotyping was performed using standard procedures. Male WT C57BL/6J (Jackson Laboratory, Bar Harbor, ME) and ESRP2-KO^20^ mice were fed an extended version (20 days + 2 binges) of the NIAAA model ^62^. Briefly, both WT (n=8) and *Esrp2*-KO mice (n=8) were initially fed the control Lieber-DeCarli diet *ad libitum* for 5 days to acclimatize them to the liquid diet. Subsequently, ethanol-fed groups were allowed free access for 20 days to the ethanol diet containing 5% (v/v) ethanol, and control groups were pair-fed with the isocaloric control diet. At days 10 and 20, ethanol-fed and pair-fed mice received a dose of ethanol (5 g/kg body weight) or isocaloric maltose dextrin, respectively, via gavage in the early morning and were euthanized 24 hours after the second binge. Whole liver tissues and hepatocytes were isolated from mice following guidelines for euthanization and/or anesthesia. Paraffin-embedded liver sections were stained with hematoxylin and eosin (H&E) or Sirius Red for pathological evaluation, and optimal cutting temperature (OCT)-embedded liver sections were stained with Oil Red O for visualization of lipid droplets. For immunohistochemistry, liver sections were deparaffinized, hydrated and incubated in 3% hydrogen peroxide to block endogenous peroxidase. Antigen retrieval was performed by heating in 10 mM sodium citrate buffer (pH 6.0) for 10 min using a microwave. Specimens were blocked in Dako Protein Block solution (Agilent, Santa Clara, CA) for 30 min at room temperature followed by incubation with primary antibody at 4 °C overnight. Other sections were also incubated at 4 °C overnight in non-immune sera. Polymer-horseradish peroxidase (HRP) anti-rabbit (Dako) and polymer-HRP anti-goat (ThermoFisher Scientific, Waltham, MA) were used as secondary antibodies and 3,3′-diaminobenzidine (DAB) as brown color was used to visualize the protein. Sections were counterstained with hematoxylin.

### Nuclei isolation and multi-omics library preparation

Intact nuclei were isolated from human liver samples based on a protocol adapted from 10X genomics CG000375. Briefly, frozen tissue was transferred to a pre-cooled Dounce homogenizer with 1x Homogenization Buffer described in ^63^ (supplemented with 0.5 U/ul RNaseIn and 1:100 dilution of PIC III). The sample was homogenized 15x using a Pellet Pestle on ice and incubated for 6 minutes on ice. The sample was homogenized again 5x using a Pestle on ice and incubated for 4 more minutes on ice. Then, the sample was passed through a suspension through a 70 μm strainer and then through a 15 μm strainer and centrifuged at 500 rcf for 5 min at 4°C. Pellet was left with 1 ml PBS + 1% BSA + 1U/μl RNase Inhibitor for 5 minutes on ice without mixing and then resuspended. Finally, this nuclear suspension was centrifuged at 500 g for 5 mins at 4°C and the obtained pellet was resuspended in 1ml PBS + 1% BSA + 1U/μl RNase Inhibitor and used for library preparation.

### Single-nuclei library preparation and sequencing

Following nuclear isolation, single-nuclei sequencing libraries for multi-omic profiling using scRNA-seq and scATAC-seq were prepared individually from pools of human liver samples from Normal individuals, AH patients and AC patients using the 10x Genomics Chromium Single Cell Kit and sequenced with Illumina NovaSeq 6000 on a SP/S4 flow cell to obtain 150bp paired reads. For each condition, we pooled the nuclei into a batch of 3 individuals and another batch of 2 individuals during library preparation and targeted 9000 and 6000 single nuclei per batch respectively.

### snRNA-seq/snATAC-seq data analysis

Raw data obtained from sequencing were demultiplexed, filtered, and mapped to the hg38 human genome using the Cell Ranger ARC pipeline. Cell Ranger ARC was also used to perform barcode counting, peak calling, counting of ATAC and GEX modules, and generating feature-barcode matrices. The data were integrated using the Seurat pipeline and batch-corrected using the harmony batch correction protocol^64^. Finally, Seurat^65,66^ and Signac^67^ pipelines were used to perform dimensionality reduction, cluster determination, cell type annotation and perform differential analysis on clusters.

### RNA and protein analysis

RNAs were extracted using TRIzol reagent according to standard protocols. Real-time PCR reactions were performed in triplicate or duplicate using the SYBR green master mix. For checking inclusion levels of exons, RT-PCR assays were done with primers spanning flanking exons and products were resolved on a 5% acrylamide gel. For protein analysis, frozen human liver samples were homogenized in RIPA lysis buffer. Protein lysates were cleared by spinning the samples twice at 4°C. Subsequently, samples were separated on SDS-PAGE and analyzed by western blotting.

### Illumina RNA-seq

Frozen tissue was quickly thawed and disrupted on ice by homogenizer. Total RNA was isolated using a Qiagen RNeasy© RNA isolation kit. After RNA quality had been assessed by capillary electrophoresis (bioanalyzer), cDNA libraries were prepared using TruSeq© RNA Library Prep Kit and analyzed with an Illumina NextSeq500. Base-calling and fastq conversion was performed using RTA (2.4.11) and Bcl2fastq (2.18.0.12), respectively.

### RNA-seq analysis

For analysis of RNA-seq data, raw reads were subjected to read length and quality filtering using Trimmomatic V0.38 ^68^ and aligned to the mouse genome (mm10) using STAR (version 2.6.1d)^69^. Cufflinks package ^70^ was used to assess differential gene expression events, among which significant events were identified using a stringent cutoff criteria (FDR(q-value) < 0.05, log2(fold change) > 1). rMATS v4.0.2(turbo) ^71^ was used to study differential splicing and events with FDR < 0.1, junction read counts ≥ 10, PSI ≥ 10% were deemed to be significant.

### Exon ontology analysis

Exon ontology analysis was performed on the set of alternatively spliced cassette exons identified using rMATS. Mouse(mm10) annotations were converted to human (hg19) annotations using UCSC liftover with a minimum base remap ratio set to 0.8. Exon ontology pipeline ^43^ was then used on the lifted exons to perform ontology analysis.

### eCLIP library preparation & analysis

ESRP2 eCLIP was performed in concordance with previously published protocols^72^. Isolated hepatocytes from N-terminal FLAG-tagged ESRP2 mice (*ESRP2^Tg/wt^*) mice were suspended in 1x PBS and crosslinked with 1 pulse of 400 mJ/cm^2^ of 254 nm UV radiation to stabilize RBP– RNA interactions. Subsequent immunoprecipitation (IP) of FLAG ESRP2-RNA complexes, RNA isolation, library preparation and sequencing were performed as described previously^73^. For pulldowns, FLAG M2 antibody (Sigma Aldrich, F1804) was pre-coupled to sheep anti-rabbit IgG Dynabeads (Thermo Fisher 11203D). Libraries were sequenced on the NovaSeq6000 platform (Illumina). eCLIP was performed on IP from two independent samples along with paired size-matched input before the IP washes.

To analyze the eCLIP reads, the CLIP tool kit (CTK) pipeline was used^74^. Briefly, 3’ adaptors were clipped from reads, followed by collapsing of PCR duplicates and removal of the N10 random barcode sequence. Reads were then mapped using the bwa aligner and further cleaned up for PCR duplicates and alignments to repetitive and ribosomal non-coding RNA regions. Peak calling was performed on the remaining mapped reads as well crosslinking site analysis was performed using CIMS/CITS.

### Cell culture

AML12 cells (ATCC® CRL-2254™) were seeded into 6-well plates (at a density of 2.0 × 10^5^ cells per well). Sterile cover glasses were placed in the 24-well plates (at a density of 1.0 × 10^5^ cells per well) and cultured in DMEM/F_12_ medium supplemented with insulin–transferrin–selenium (cat. no. 41400-045; Invitrogen) and 10% fetal bovine serum (heat-inactivated at 56 °C) at 37 °C in the presence of 95% air and 5% CO_2_. For immunofluorescence, AML12 cells were seeded on sterile cover glasses placed in the 6-well plates. At 70-80% confluency, growth media was replaced with low serum media (2% FBS) and incubated for 4h prior to TGF-β treatment. Using a final concentration of 5 ng/mL TGF-β (cat. no. 7666-MB; R&D Systems), cells were incubated for 36h before harvesting. For recovery experiments, the cells were treated with 15 μM TGF-β inhibitor (LY2109761, cat. no. HY-12075; MedChemExpress) for 36h-48h. For activating beta-catenin, AML12 cells at 70% confluency were treated with 3 μM CHIR99021 (cat. no. 4423; Tocris) for 48h. To induce Tcf4 73-nt skipping, an antisense morpholino oligomer (Tcf4_73 ASO) sequence was used: 5’-ATCTGGGAGACTATACACACCTGCA-3’, which binds to the 5’ss downstream of Tcf4 exon 73 pre-mRNA sequence. As a control, a non-target standard control sequence was used: 5’-CCTCTTACCTCAGTTACAATTTATA-3’ (Gene Tools, LLC, Philomath, Oregon, USA). Morpholinos were delivered at 10uM final concentration using Endo-porter reagent 48h. Cells were harvested to prepare total RNA and protein lysates.

### Immunofluorescence staining and analysis

AML12 cells with control and experimental treatments were grown on coverslips and harvested between 24h-48h, fixed in 4% paraformaldehyde for 10 minutes, permeabilized using 0.5% Trition-X100 in 1X PBS for 5 minutes, blocked in 2% Normal Goat Serum, 1% BSA, 0.1% Triton X100 in 1X PBS, and incubated with primary antibody (1:1000) overnight at 4°C in a humidified chamber. On the following day, the cells were incubated with a secondary antibody (1:500) at room temperature for 1-2h, treated with NucBlue for 20 minutes, and mounted. All the images were taken using Zeiss LSM 900 with Airyscan 2 at the core facilities of IGB, UIUC. The images were processed using Airyscan Deconvolution and ImageJ, and >10 nuclei per sample were randomly selected for analysis across biological replicates. These images were analyzed using our custom CellProfiler pipeline, wherein nuclei were defined and separated from the surrounding non-nuclear region using Otsu’s global thresholding method. Intensity for SLK and TCF4 were separately measured for the nuclear and non-nuclear regions and the nuclear fraction was calculated as a ratio of nuclear to non-nuclear regions. The data was analyzed in Prism using unpaired non-parametric Mann-Whitney U Test (obtained p-value <0.0001).

### RNA-seq Data Normalization and Dimensionality Reduction

We normalized the RNA-seq dataset using the Seurat pipeline^28^. This process involved the following steps: NormalizeData(object, Normalization.method = “LogNormalize”), FindVariableFeatures (object, selection.method = “vst”, nfeatures = 2000, clip.max = “auto”, binning.method = “equal_width”), ScaleData(model.use = “linear”, min.cells.to.block = 3000, block.size = 1000, scale.max= 10), and RunPCA(npcs = 50, weight.by.var = TRUE). The resulting normalized “RNA” assay was used for subsequent analyses.

### Porto-Central Coordinates Calculation

To determine the position of each cell along the porto-central axis of the hepatic lobule, we calculated a porto-central coordinate (η) based on the expression of previously identified portal and central marker genes in mice^34^. It’s important to note that while these marker genes were identified in mouse samples, we applied them to our human RNA-seq dataset, assuming the conservation of zonation patterns across species.

Portal marker genes (*pLM*) included: *APOF, APOM, ASGR2, ASS1, C1S, C8B, CPT2, EEF1B2, ELOVL2, FADS1, FBP1, GC, GNMT, HSD17B13, IFITM3, IGF1, IGFALS, NDUFB10, PIGR, S100A1, SERPIND1, SERPINF1, UQCRH, VTN, PCK1, ARG1*, and *CPS1*.

Central marker genes (*cLM*) included: *ALAD, ALDH1A1, C6, CPOX, CSAD, CYP1A2, CYP2E1, GSTM1, HPD, LECT2, MGST1, OAT, PON1, PRODH, RGN,* and *SLC16A10*.

For each cell, *i*, we calculated the identified porto-central coordinate *η_i_* using the following equations:

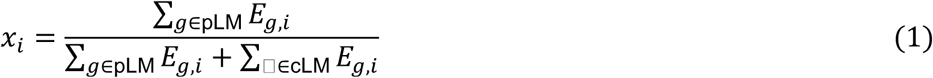

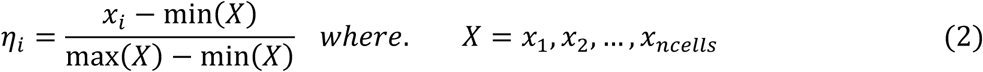

Where *E_g,i_* represents the expression of gene, *g*, in cell, *i*.

The resulting *η_i_* values range from 0 to 1, with *η_i_* = 0 indicating a position closer to the pericentral region and *η_i_* = 1 indicating a position closer to the periportal region of the hepatic lobule. It should be noted that some genes from the original list (*CML2, CYB5, CYP2C37, CYP2C50, CYP3A11, ATP5A1, CYP2F2, DAK, SERPINA1C, SERPINA1E,* and *TRF*) were not found in our dataset and were therefore excluded from the analysis.

### Zonation and Comparative Analysis

We divided the hepatic lobule into three distinct zones based on its calculated porto-central coordinate *η_i_*. Given that *η_i_* ranges from 0 to 1, we equally partitioned this range into three zones:

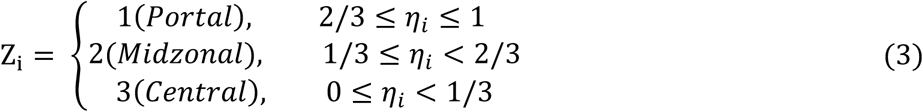

Where *Z_i_* represents the zone assignment and *η_i_* is the porto-central coordinate of each cell, *i*. We compared the zonation patterns between different hepatic conditions (Normal, Alcohol-associated Cirrhosis [AC], and Severe Alcohol-associated Hepatitis [SAH]), we implemented a subsampling and distance calculation approach using R (R Core Team, 2021) and the Seurat package^28^. We performed two rounds of random subsampling without replacement. The number of cells subsampled was determined by the minimum cell count across conditions in the same zone, ensuring equal representation. The maximum amount of cells we subsampled across conditions is 100 to reduce noise and ensure a proper representation. This process resulted in two independent subsets of cells for each zone-condition combination, allowing for robust comparative analysis.

Using the principal component (PC) embeddings from the Seurat objects, for any two cells *i* and *j* in the same zone in the subsample, we computed the Euclidean distance *d_ij_* in the n-dimensional PC space (Equation 4).

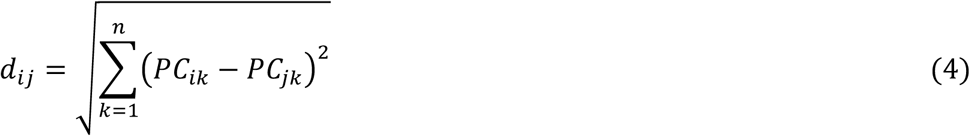

Where *PC_ik_* and *PC_ik_* represent the kth principal component values for cells *i* and *j*. This calculation provides a quantitative measure of cell-to-cell variability in the reduced dimensional space.

### Zonation-based Gene Expression Analysis

We developed a method inspired by the Methods of the Halpern et al. paper to investigate gene expression patterns across hepatic zones in different conditions (Normal, AC and SAH)^34^. The set of codes created generates three key data structures: the Porto-central Matrix (*M* ∈ *R*^#*cells*,#*zones*^), Mean Gene Expression (MGE) matrix and a variance-sorted gene list.

Using the zonation data obtained from the previous section, we constructed a binary matrix called the Porto-central Matrix (*M* ∈ *R*^#*cells*,#*zones*^), where rows represent cells *i* and columns represent zones *Z_i_*. Each cell was assigned a value of 1 in its corresponding zone column and 0 in others. To ensure balanced representation across zones, we subsampled an equal number of cells (n = 28) from each zone, resulting in a subsampled matrix. The matrix is then normalized through a two-step process: first, each row is divided by its sum, and then each column is divided by its sum. This can be expressed mathematically as:

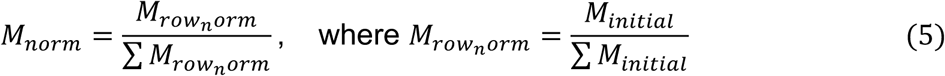

Using this weighted normalized matrix and the gene expression data from our Seurat objects (*E* ∈ *R*^#*genes*,#*column*^), we obtain the Mean Gene Expression matrix (*G* ∈ *R*^#*gen*,#*zones*^) with rows of genes and columns of normalized average expression of each gene across zones (Equation 6).

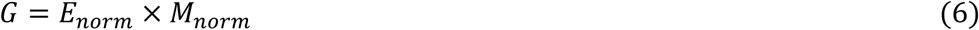

We then calculated the variance of gene expression for each gene in the normalized Mean Gene Expression matrix *G* (Equation 7).

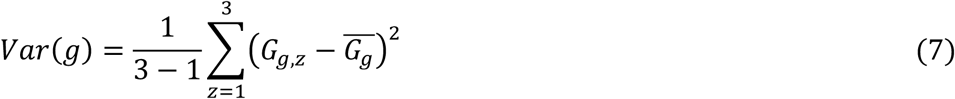

where *G_gZ_* represents the expression of gene *g* in zone *Z*, and *G̅_g_* is the mean expression of gene *g* across all three zones. We then ranked genes in descending order based on their calculated variances. To focus on identifying novel zonation-associated genes, we excluded the porto-central gene markers used in the initial zonation calculation from this ranked list.

For any two zones *Z_i_* and *Z_j_* in the condition, we computed the Euclidean distance *d_ij_* in the n-dimensional gene space (Equation 8). This computation uses only genes with a variance of 0.1 and above for each zone. This analysis pipeline was applied separately to each condition (Normal, AC, and SAH).

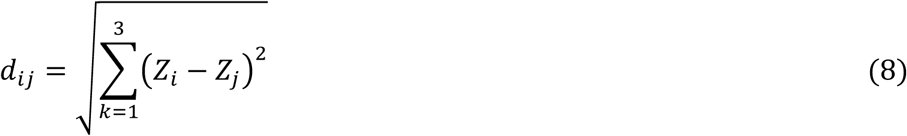

### Zonation Mean Gene Expression Analysis

We extracted normalized gene expression data from Seurat objects for Normal liver and Severe Alcohol-associated Hepatitis (SAH) conditions. The data was subset based on a predefined list of zonation genes, including both portal and central markers, compiled from previous liver zonation literature.

For each condition (Normal and SAH) and zone (1, 2, and 3), we calculated two types of mean gene expression for each zonation gene: 1) the mean expression across all cells, including those with zero expression, and 2) the non-zero mean expression, focusing only on cells where the gene is detected. This dual approach accounts for both overall expression levels and the sparse nature of single-cell RNA-seq data. The results were compiled into two separate matrices of mean gene expression, each with rows representing individual zonation genes and columns representing each zone within Normal and SAH conditions (6 columns total). For both matrices, genes not detected in any cells for a particular zone and condition were marked as NA. We included genes in the analysis if they showed expression in at least one cell in each zone and condition.

### Statistical analysis

We employed two complementary statistical tests: the Wilcoxon rank-sum test and the independent two-sided t-test. Cells in each zone in different conditions are independent of each other, we conducted both tests to compare the significance of differences in the distribution of cells between conditions within zones and in gene expression between zones in different conditions. The Wilcoxon test, being non-parametric, allowed us to compare the distribution of cells within zones and the value differences in gene expression between zones without assuming normality, while the t-test provided insights into differences in the mean distribution of cells and expression levels. We considered p-values < 0.05 as statistically significant for both tests.

## Code availability

All code developed for portal-central coordinate calculation, zonation characterization, statistical analysis, variability of gene expression calculation, and plotting can be found on GitHub at https://github.com/GoyalLab/Zonation_Analysis

## SUPPLEMENTAL INFORMATION

Supplemental information can be found online at

## ACKNOWLEDGMENTS

We acknowledge support from the Transgenic mouse core, High-throughput sequencing and genotyping core, and Histology-microscopy core facilities at the University of Illinois, Urbana-Champaign. We thank the members of the Kalsotra and Diehl laboratories for their valuable discussions and comments on the manuscript. Figures were created with BioRender.com.

## AUTHOR CONTRIBUTIONS STATEMENT

UVC and SB designed the research studies, conducted experiments, acquired data, performed computational analysis, analyzed data, wrote and revised the manuscript; RD, DD, SN, KT, AB and IP conducted experiments and acquired data; ZS and BP provided human liver samples and bulk RNA-seq data; AL and YG performed computational analysis, AK and AMD designed the research studies, analyzed data, wrote and revised the manuscript, secured funding for the research. All authors discussed the results and edited the manuscript.

## SUPPLEMENTAL FIGURE LEGENDS

**Supplementary Figure 1:**
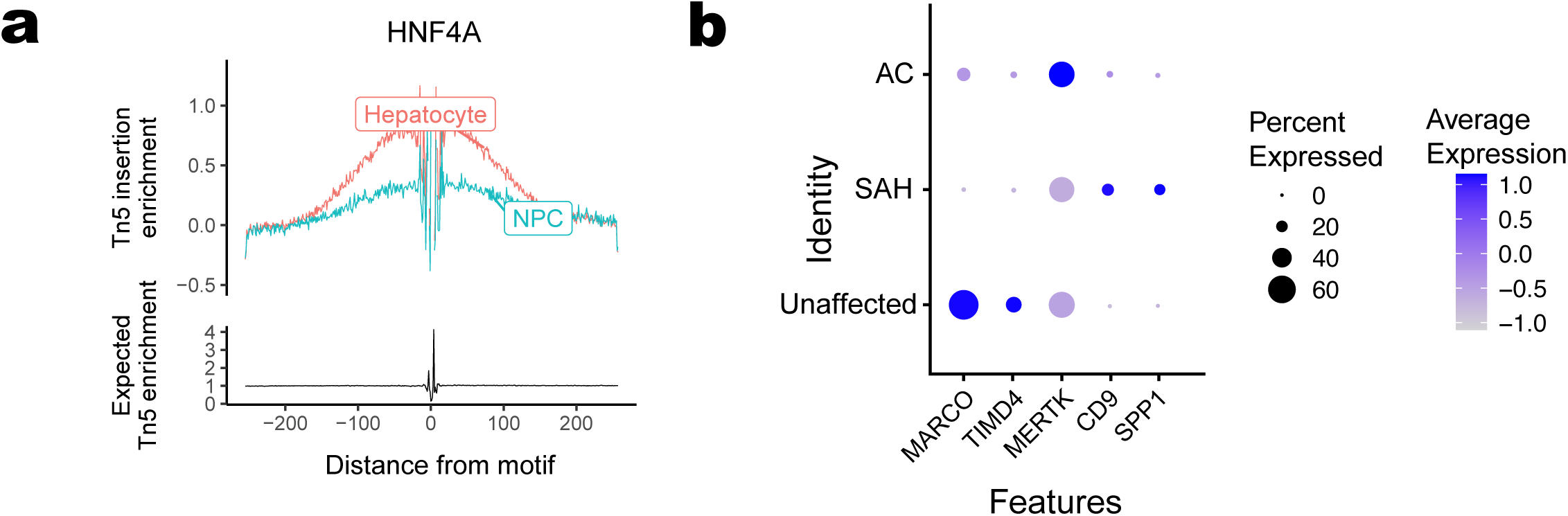
**(a)** Genome-wide Tn5 insertion enrichment plots around motifs for HNF4α indicating their increased accessibility in hepatocytes compared to hepatocytes **(b)** Dot plot showing disease-dependent changes in expression levels of markers for Kupffer cell (MACRO, TIMD4, MERTK) and scar-associated macrophages (TREM2, CD9, SPP1).

**Supplementary Figure 2:**
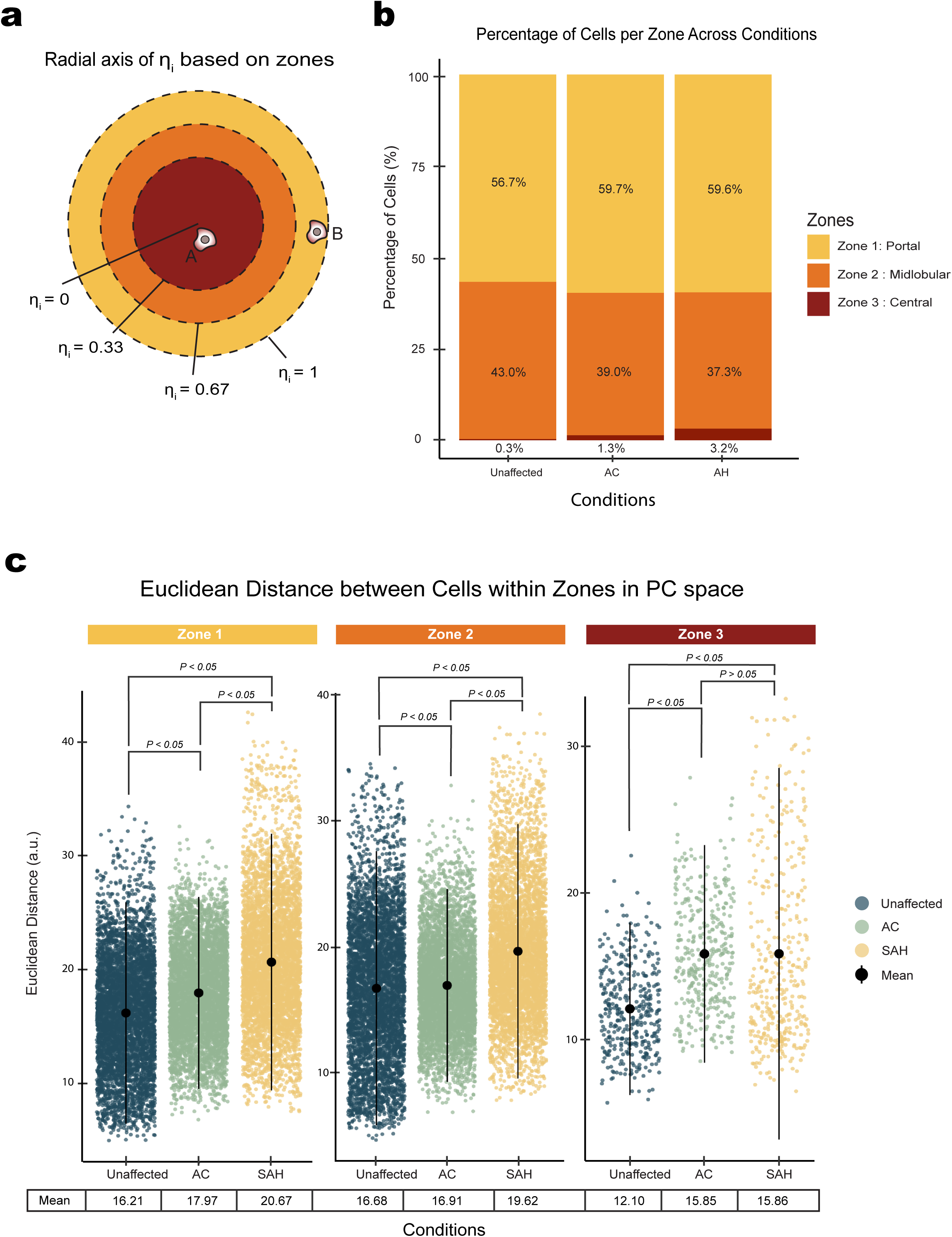
**(a)** Schematic representation of the radial axis of *η_i_* based on hepatic lobule zones. The concentric circles illustrate the equal division of the lobule into three zones, with *η_i_* values ranging from 0 (central) to 1 (portal). Cell A is closer to the central axis and Cell B is closer to the portal axis. **(b)** Stacked bar chart displaying the percentage of cells per zone across Unaffected, SAH, and AC conditions. Zone 1 represents the Portal Axis and Zone 3 represents the Central Axis. **(c)** Euclidean distances between cells within hepatic zones in principal component (PC) space. Scatter plot showing pairwise Euclidean distances between cells within each zone (1, 2, 3) for Unaffected, SAH, and AC conditions.

**Supplementary Figure 3:**
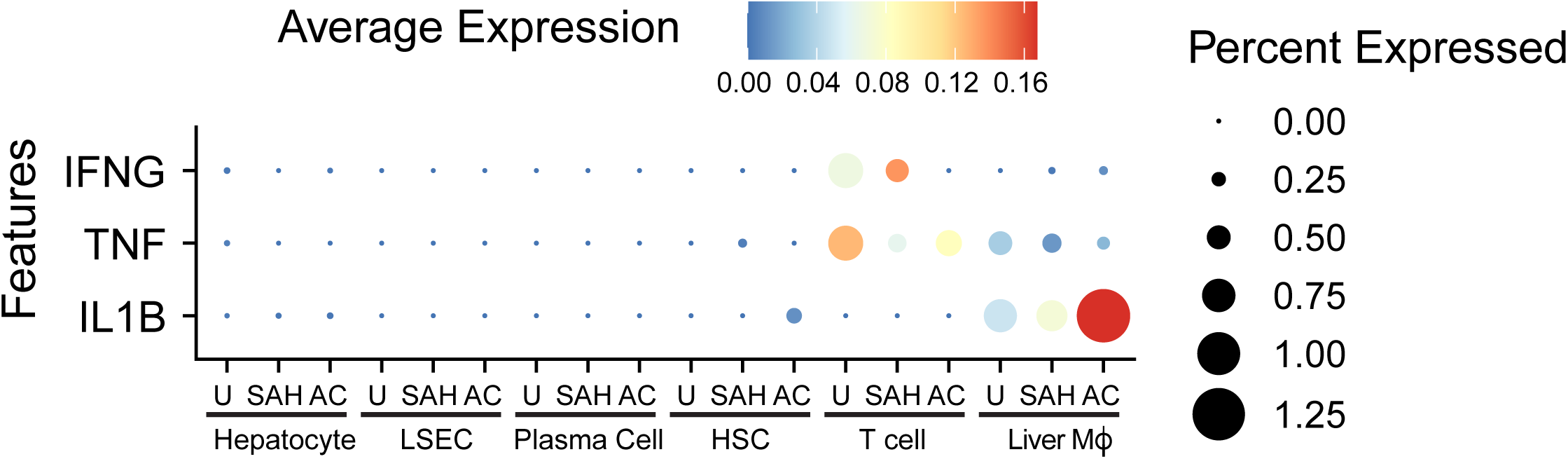
Cell type-specific expression levels of key pro-inflammatory cytokines from snRNA-seq dataset.

**Supplementary Figure 4:**
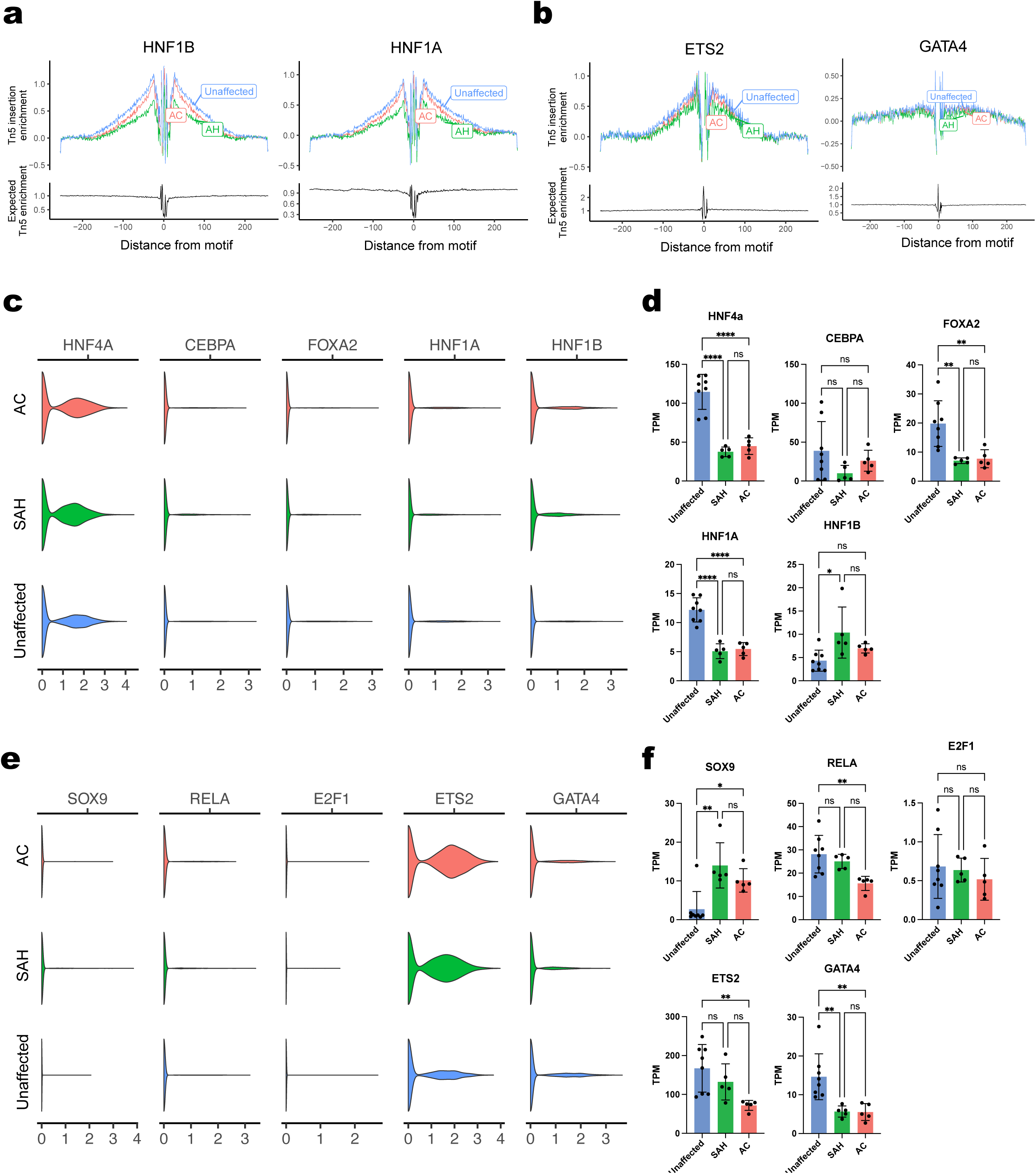
Genome-wide Tn5 insertion enrichment plots for **(a)** HNF1A and HNF1B, **(b)** ETS2 and GATA4, comparing activities across unaffected, SAH and AC livers. Expression levels of transcription factors active in adult hepatocytes from **(c)** multi-omics dataset and **(d)** bulk RNA-seq dataset. Expression levels of transcription factors more active in fetal livers from **(e)** multi-omics dataset and **(f)** bulk RNA-seq dataset.

**Supplementary Figure 5:**
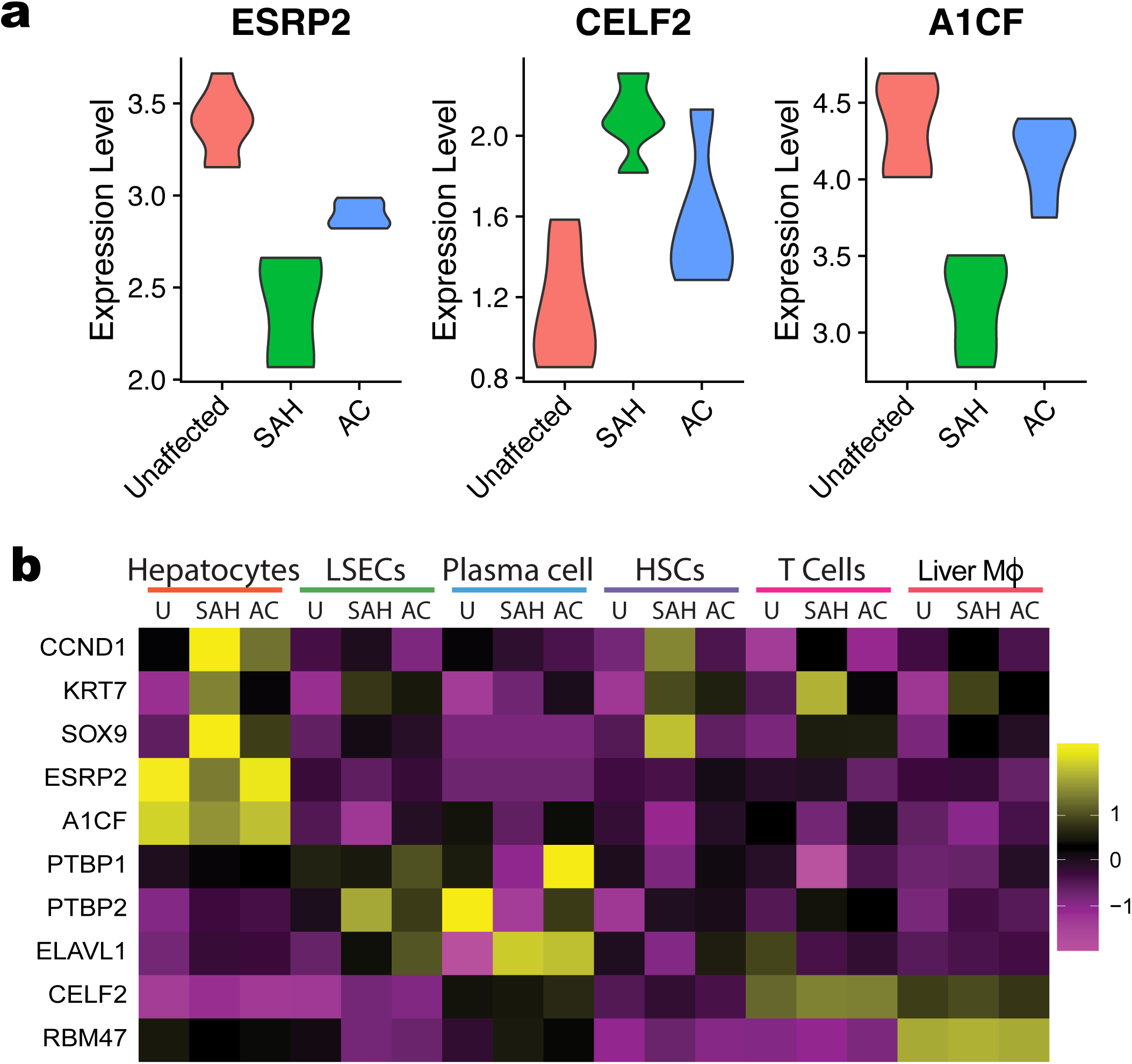
**(a)** Violin Plot showing expression of ESRP2 and CELF2 across liver pathologies from bulk RNA sequencing data. **(b)** Heatmap from snRNA-seq module showing cell type-specific changes in expression of progenitor/fetal-like markers and RNA binding proteins from alcohol-associated liver disease (ALD).

**Supplementary Figure 6:**
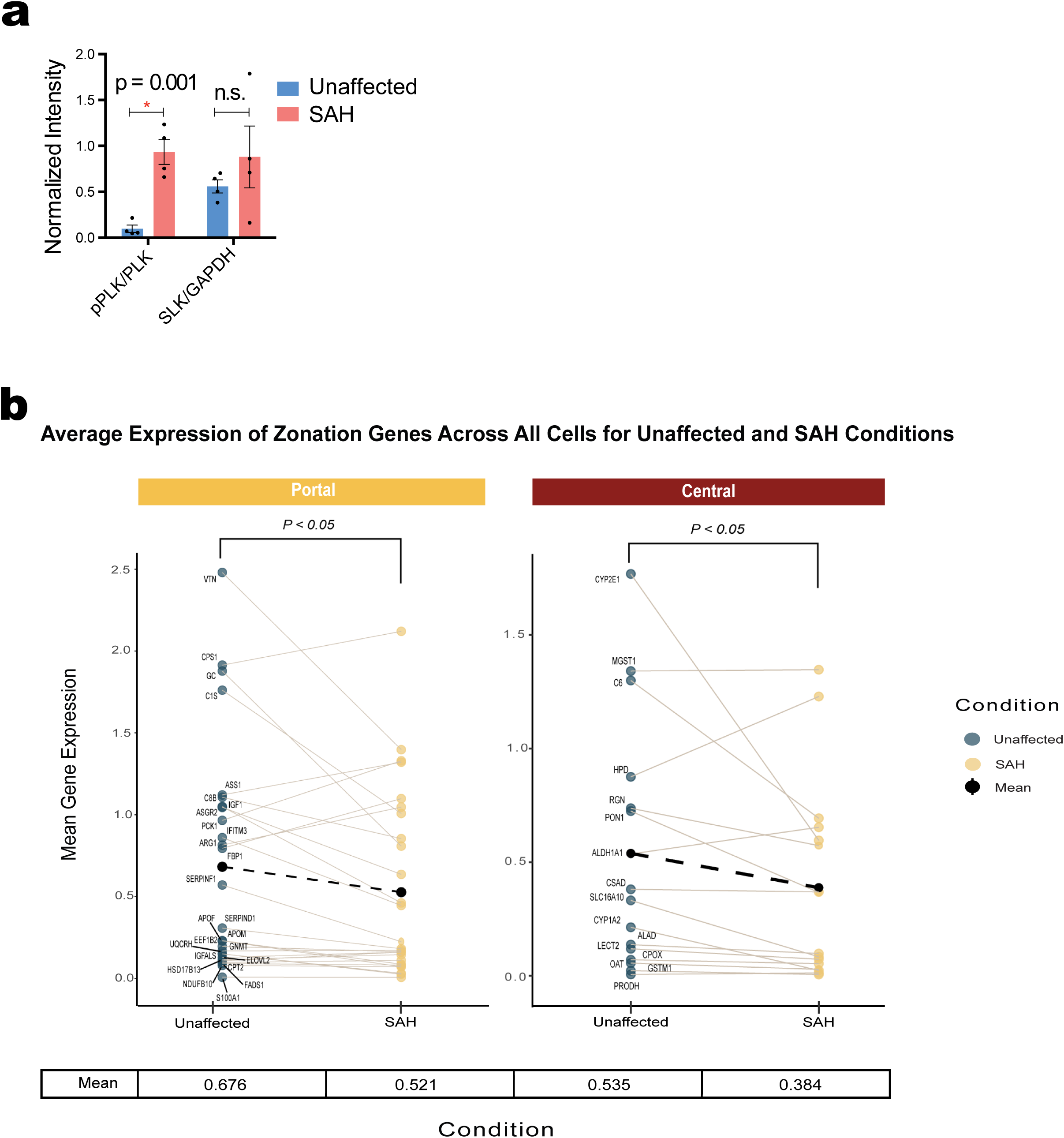
**(a)** Quantitation for western blot analysis from human liver lysates demonstrating increased phosphorylation of Polo-like kinase (PLK) in SAH patients. GAPDH blot shown in Figure 3E is used for normalization. **(b)** Plots showing average expression of zonation Genes across all cells for unaffected and SAH conditions.

**Supplementary Figure 7:**
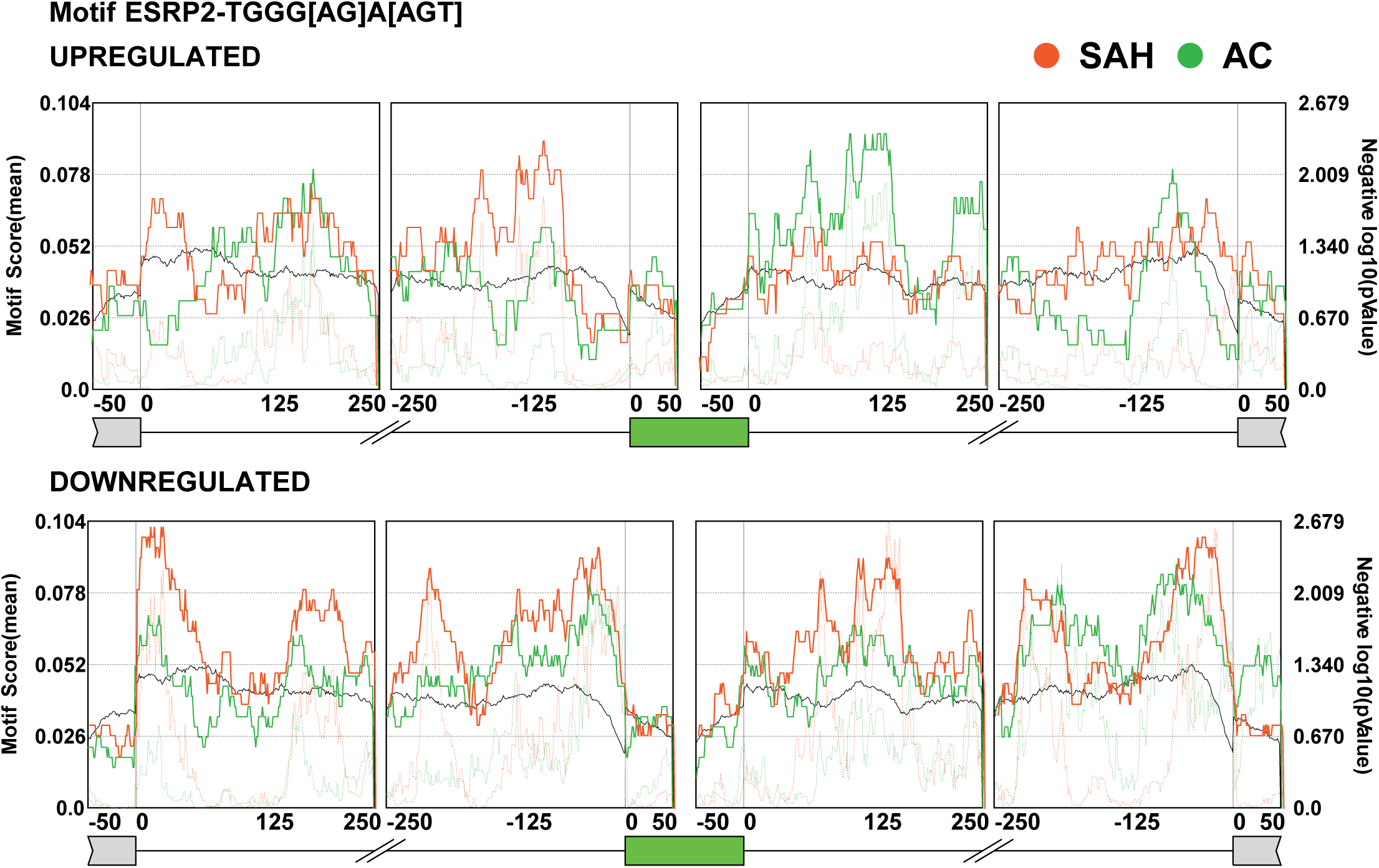
Enrichment plots for ESRP2 binding motif presence in alternative exons and surrounding intronic regions that are misregulated in SAH and AC.

**Supplementary Figure 8:**
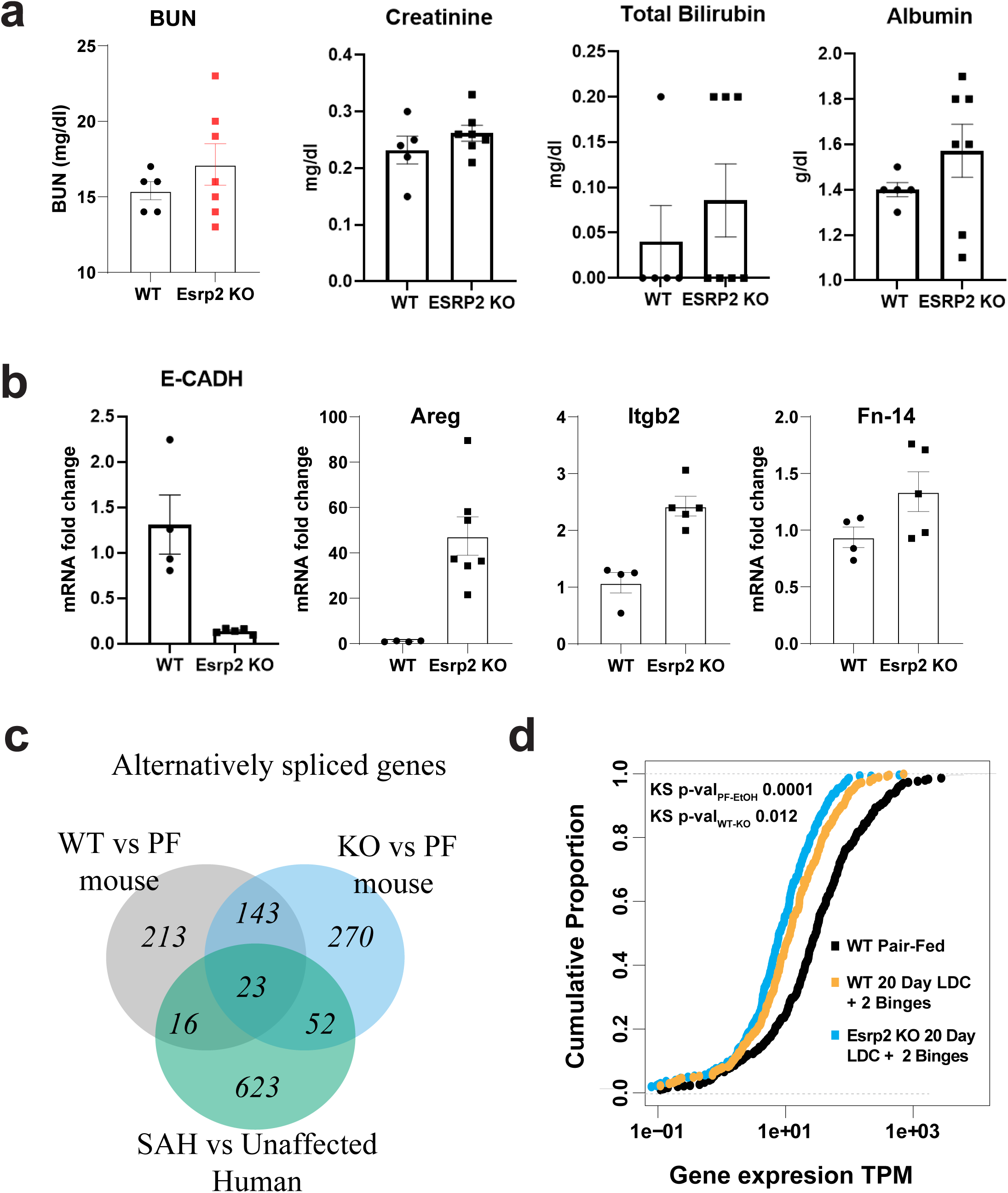
**(a)** Serum levels of BUN, Creatine, total bilirubin and Albumin in ESRP2 KO as compared to control mice after extended NIAAA diet schedule (20D+2B). **(b)** qRT-PCR based quantitation of selected genes from ESRP2 KO mice liver relative to WT C57BL6 mice, after extended NIAAA diet schedule. **(c)** Venn diagram showing overlap of genes that are alternatively spliced in WT and ESRP2 KO mice upon 20D+2B diet relative to pair-fed WT control mice. **(d)** Cumulative plot illustrating trends in expression of TCF4 target genes in 20D+2B fed WT and ESRP2 KO mice liver in comparison to pair-fed WT control mice liver.

